# Rapid Reconfiguration of the Functional Connectome after Chemogenetic Locus Coeruleus Activation

**DOI:** 10.1101/527457

**Authors:** Valerio Zerbi, Amalia Floriou-Servou, Marija Markicevic, Yannick Vermeiren, Oliver Sturman, Mattia Privitera, Lukas von Ziegler, Kim David Ferrari, Bruno Weber, Peter Paul De Deyn, Nici Wenderoth, Johannes Bohacek

**Author notes:** **equal contribution** (Valerio Zerbi, Amalia Floriou-Servou). **corresponding authors:** Valerio Zerbi, Nicole Wenderoth, Johannes Bohacek.

## Abstract

The locus coeruleus (LC) supplies norepinephrine (NE) to the entire forebrain, regulates many fundamental brain functions, and is implicated in several neuropsychiatric diseases. Although selective manipulation of the LC is not possible in humans, studies have suggested that strong LC activation might shift network connectivity to favor salience processing. To test this hypothesis, we use a mouse model to study the impact of LC stimulation on large-scale functional connectivity by combining chemogenetic activation of the LC with resting-state fMRI, an approach we term “chemo-connectomics”. LC activation rapidly interrupts ongoing behavior and strongly increases brain-wide connectivity, with the most profound effects in the salience and amygdala networks. We reveal a direct correlation between functional connectivity changes and transcript levels of *alpha-1, alpha-2*, and *beta-1* adrenoceptors across the brain, and a positive correlation between NE turnover and functional connectivity within select brain regions. These results represent the first brain-wide functional connectivity mapping in response to LC activation, and demonstrate a causal link between receptor expression, brain states and functionally connected large-scale networks at rest. We propose that these changes in large-scale network connectivity are critical for optimizing neural processing in the context of increased vigilance and threat detection.

## INTRODUCTION

The locus coeruleus (LC) is a small structure in the brainstem (with approximately 1500 neurons in each hemisphere in mice and 20’000 in humans)^1,2^, which sends widespread efferent projections to almost the entire brain, and constitutes the major source of norepinephrine (NE) to most forebrain regions. Dysregulation of the LC-NE system has been implicated in numerous psychiatric pathologies including depression, anxiety, attention-deficit-hyperactivity disorder, post-traumatic stress disorder and neurodegenerative diseases^3–8^. The ability to selectively change activity within the LC-NE system with optogenetic and chemogenetic manipulations has confirmed that the LC has a strong modulatory effect on various functional circuits related to wakefulness^9^, cognitive performance and memory^6,10–13^, and on stress-related behavioral responses including fear, anxiety and avoidance^3,5,14–19^. These widespread functional effects are in line with theories that the LC optimizes cognitive processes relevant for task performance or adaptive behaviors by rearranging neural activity within and between large-scale neuronal systems^1,20–23^.

Phasic LC activation, as triggered by salient stimuli, enhances cognitive performance and facilitates faster orienting towards (and processing of) task-relevant cues^1,20,24^. In contrast, high tonic LC activity, as reliably and robustly triggered by various stressors, causes the release of substantial quantities of NE throughout the brain^25,26^, which is thought to have a “circuit breaker” function that allows the interruption of ongoing neural activity and rapid reconfiguration of functional communication between brain regions (i.e. functional networks)^20,26,27^. This rapid response is highly evolutionarily conserved, as it benefits survival by enhancing vigilance and enabling the selection of adaptive behaviors in threatening situations^28^. However, it has not been demonstrated directly that increased LC activity reconfigures functional neural networks across the brain, and it remains unknown how the widespread LC projections might achieve specificity for regulating specific networks. Some evidence suggests that strong LC activation by environmental cues, as observed during acutely stressful situations, plays an important role in activating networks that favor salience processing and action selection^20,29^, and that regional specificity is achieved through the distribution of adrenergic receptors^26^. In humans, acute stress exposure dynamically shifts large-scale network activity towards higher activation of the salience network (including amygdala)^30^, which is mediated by beta-adrenergic receptors^31^ and is thought to promote hypervigilance and threat detection at the cost of executive control^27,29^. However, direct involvement of the LC has not been proven, since it is impossible to selectively manipulate LC activity in humans.

To explore whether LC activation changes local and global network organization, we used a translational mouse model and applied a novel ‘chemo-connectomics’ approach, which combines (i) cell-specific chemogenetic manipulation of neural activity afforded by Designer Receptors Exclusively Activated by Designer Drugs (DREADDs)^32^; with (ii) a brain-wide functional connectome analysis as revealed by resting state functional magnetic resonance imaging (rs-fMRI) in mice. This approach leverages the molecular tools and genome-wide resources available in mice and allows us to link functional connectivity with micro- and mesoscopic properties of the mouse brain. We asked (1) whether a selective increase of LC activity would change the functional connectivity profiling of large-scale brain networks or “connectome”, (2) if such network-wide effects are related to the known distribution of adrenergic receptor subtypes across the brain, and (3) whether changes in functional connectivity correspond to the levels of NE release in the target structures.

## RESULTS

To selectively target the LC, we used transgenic mice that express codon-improved Cre-recombinase (iCre) under the dopamine-*beta*-hydroxylase (DBH) promoter (DBH-iCre mice, Figure 1A)^33^. We stereotactically delivered floxed excitatory DREADDs^34^ (AAV5-hSyn-DIO-hM3Dq-mCherry; *hM3Dq-mCh*) or a control virus (AAV5-hSyn-DIO-mCherry; *mCh*) to the LC, thus restricting virus expression to DBH-positive noradrenergic neurons of the LC (Figure 1B). We assessed successful LC activation using pupillometry, a highly sensitive and clinically relevant readout of LC activation^35–37^. After two minutes of baseline recording under light isoflurane anesthesia, we activated LC neurons by administering the potent DREADD activator clozapine at an ultra-low dose (0.03 mg/kg, i.p., Figure 1C)^38^. Within a minute of clozapine injection, we observed a strong increase in pupil diameter in the hM3Dq-mCh group, while pupil diameter of mCh mice did not change in response to clozapine injection and remained stable throughout the 10-minute recording session (Figure 1D-F). To show that our LC activation protocol is behaviorally relevant, we subjected mice to an open field test (OFT) immediately after clozapine injection and recorded their behavior for 30 minutes. In comparison to mCh mice, clozapine injection had profound effects on the behavior of hM3Dq-mCh mice. Several minutes after clozapine administration, hM3Dq-mCh mice showed strongly suppressed locomotor activity (Figure 1G, H), spent less time in the (more aversive) center of the open field (Figure 1I, J), performed less activity-related supported rears (Figure 1K, L) and less exploratory unsupported rears (Figure 1M, N)^39^. This is in line with previous findings that LC activation suppresses motor activity^9,16^ and increases anxiety^3,14,16^. Because the LC is a highly sexually dimorphic structure, we assessed whether these findings hold true in female mice as well. We confirmed that after clozapine administration, hM3Dq-mCh females also showed reduced locomotor activity in the OFT (**Figure S1B, C**), spent less time in the center (**Figure S1D, E**), and performed fewer supported (**Figure S1F, G**) and unsupported rears (**Figure S1H, I**). To better characterize the effects of LC activation on behavior, we also tested these females in the light-dark box (LDB) for 30 minutes, immediately after clozapine injection (**Figure S1A**). Compared to mCh, hM3Dq-mCh mice spent much less time in the aversive light compartment of the box (**Figure S1J, K**), and more time in the dark compartment (**Figure S1L, M**). They also travelled less distance (**Figure S1N, O**), shuttled fewer times between the light and the dark compartment (**Figure S1P**), and performed fewer rears (**Figure S1Q, R**). These results indicate reduced exploratory behavior and increased anxiety. As the strong suppression of locomotor activity could also be due to locomotor impairment, we trained the same mice on the rotarod (Day 1), and then tested them on two consecutive days, without clozapine (Day 2) or with clozapine (Day 3, see **Figure S1A**). Both during training and testing, hM3Dq-mCh mice performed similarly before and after clozapine injection (**Figure S1S**), with no significant difference between groups or across daily trials (**Figure S1T, S**1U). Thus, the strong suppression in locomotor activity after LC activation is likely due to an increase in anxiety and not due to any gross locomotor impairment.

**Figure 1.**
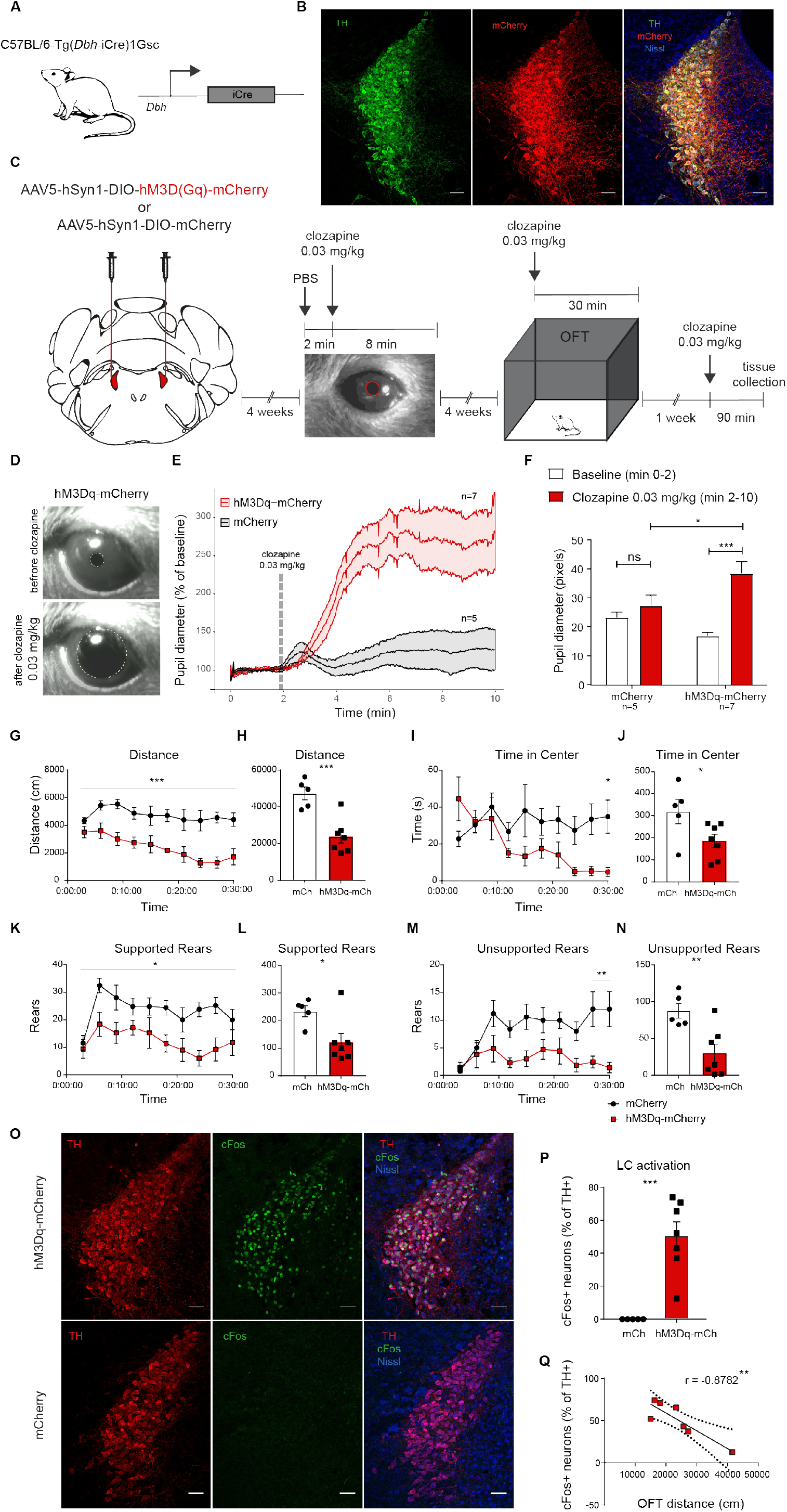
Physiological, behavioral and molecular effects of LC activation with hM3Dq. (**A**) Schematic of I *DBH*-iCre mouse genetics. (**B**) mCherry (mCh) co-localizes with tyrosine hydroxylase (TH) in the LC of DBH-iCre mice after stereo-tactic delivery of AAV5-Syn1-DIO-mCh. (**C**) Diagram of the experimental design. (**D**) Images showing the pupil size before and after administration of clozapine in a mouse expressing hM3Dq-mCh in LC. (E, F) Pupil size increased after clozapine administration only in hM3Dq-mCh mice (interaction time x group: F(1,10)=9.60, *p*=0.0113, two-way ANOVA with Sidak *post hoc* tests). (**G-N**) Immediately after clozapine injection, hM3Dq-mCh mice travelled less distance than mCh control mice (G, H: main effect of group: F(1,10)=21.92, *p*=0.0009, two-way ANOVA), spent less time in the center (I, J: main effect of group F(1,10)=5.15, *p*=0.0467; interaction: F(9,90) =3.04, *p*=0.0032, two-way ANOVA with Sidak *post hoc* tests), performed fewer supported rears (**K, L**: main effect of group: F(1,10)=7.24, *p*=0.0227) and fewer unsupported rears (**M, N**: main effect of group: F(1,10) =11.33, *p* =0.0072, interaction: F(9,90)=2.46, *p*=0.0148). (**O**) Representative images of cFos expression in TH+ neurons in the LC of hM3Dq-mCh or mCh mice, 90 minutes after injection of clozapine. (**P**) Quantification of cFos expression showing increased neuronal activation in hM3Dq-mCh mice (t(10)=5.12, *p*=0.0005, unpaired t test). (**Q**) cFos expression in LC correlates with distance travelled in the OFT in hM3Dq-mCh mice (r(5)=-0.8782, *p*=0.0093). hM3Dq-mCh (n=7), mCh (n=5). *=*p*<0.05, **=*p*<0.01, ***=*p*<0.001. Data represent mean ± SEM. All scale bars: 50 μm.

Next, we validated activation of LC neurons molecularly. To this end, we returned to the same male mice that were used for OFT testing and injected them again with clozapine, collected their brains 90 minutes later and assessed the neural activity marker cFos in the LC using immunohistochemistry. As expected, we observed a strong increase in cFos expression restricted to tyrosine hydroxylase-positive (TH+) noradrenergic neurons of the LC in hM3Dq-mCh mice, whereas cFos was virtually absent in mCh mice (Figure 1O, P). Because behavior testing and cFos staining were performed in the same mice, we correlated the locomotor activity of hM3Dq-mCh mice with the number of cFos-positive neurons in the LC. We found a strong negative correlation (Figure 1Q), showing that the strength of LC activation predicts the suppression in locomotion/exploration. Thus, our chemogenetic strategy specifically activates LC neurons and rapidly induces behavioral changes that last at least 30 minutes.

### LC activation drives rapid increases in functional connectivity

We hypothesized that an increase of LC-NE activity would rapidly reconfigure large-scale brain networks as reflected by the functional *connectome*. We therefore acquired resting state-fMRI (rs-fMRI) data before and after hM3Dq-induced LC activation in a continuous imaging session. We kept mice under light isoflurane anesthesia (see methods) and recorded 15 min of baseline fMRI, before activating hM3Dq with clozapine (0.03 mg/kg, i.v.)^40^. After the clozapine injection, we continued the fMRI recordings for 8 minutes (i.e. transitory period) followed by another 15 minutes where LC is expected to be robustly activated (i.e. active period), leading to a total uninterrupted scan time of 38 minutes (Figure 2A). Although a sustained increase in neural activity can be shown for at least two hours after hM3Dq-DREADD activation^38^, we limited the duration of the functional imaging session to reduce the accumulation of isoflurane over time, which might otherwise impact the local excitation-inhibition balance and neurovascular coupling^41^.

**Figure 2.**
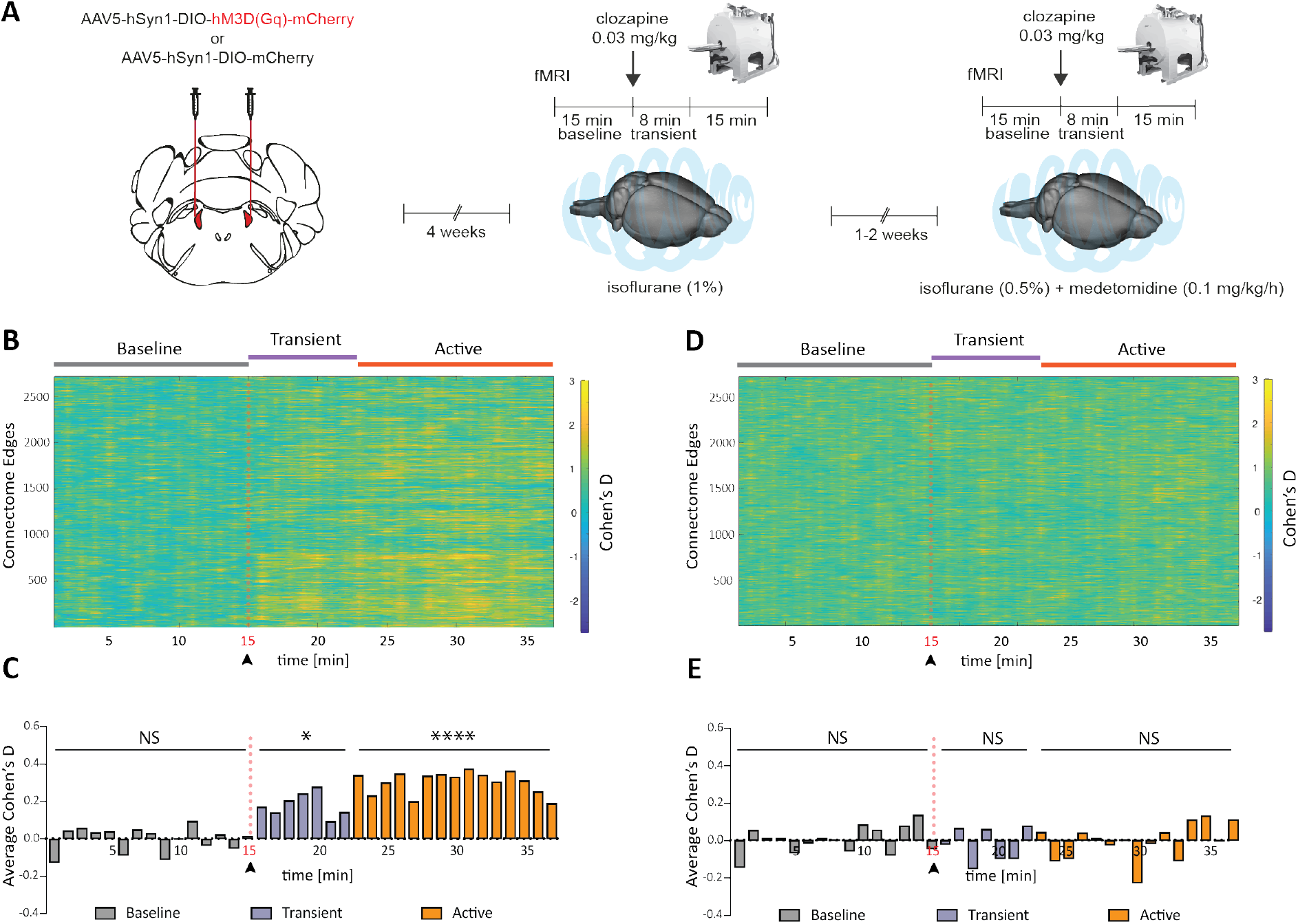
DREADD activation of the LC causes time-locked changes in functional connectivity. (**A**) Schematic of the experimental set-up for MRI recordings. 4 weeks after bilateral virus delivery, mice underwent two MRI sessions with the same experimental procedure but different anesthetic regimes. Effect-size (Cohen’s D) analysis of Functional Connectivity (FC) is shown for single-edges (n=2724) (**B**) and for the average across all edges (**C**). The data reveal time-locked increases in connectivity in multiple edges, starting immediately after clozapine (0.03 mg/kg) i.v. injection (Wilcoxon two-tailed test: *p*=0.804 for baseline period, *p*=0.015 for transient period and *p*<0.0001 for active period). (**D, E**) Treatment with the *alpha-2* adrenergic agonist medetomidine (0.05 mg/kg + 0.1mg/kg/h, see methods) abolishes clozapine-induced effects between hM3Dq-mCh and mCh groups (Wilcoxon two-tailed test: *p*=0.805 for baseline period, *p*=0.937 for transient period, andp=0.978 for active period). *=p<0.05, ****=p<0.0001, NS=not significant. hM3Dq-mCh (n=11), mCh (n=7).

We first tested whether the observed changes in whole-brain connectivity were time-locked to DREADD activation in hM3Dq-mCh mice, but not mCh mice. We measured functional connectivity (FC) between 165 brain regions using the Allen’s Common Coordinate Framework, and within 38 non-overlapping bins of 1 minute. For each bin, FC is measured between each pair of regions and normalized to the subject’s baseline connectivity (i.e. averaged across the first 15 minutes), and the effect size is calculated using the standardized difference between the group means (Cohen’s D, hM3Dq-mCh vs mCh). During the first 15 minutes (baseline) Cohen’s D varied on average between −0.12 and +0.10 (average: −0.00, null-to-small effect), and did not demonstrate an appreciable spatial or temporal pattern. However, immediately after clozapine injection, the effect size rapidly and significantly increased for the remainder of the scan session, showing increased connectivity in hM3Dq mice relative to mCh controls (Figure 2B, C). Effect sizes range on average from +0.21 to +0.38 (moderate effect) and up to +3.2 for individual edges (very strong effect). These findings show that changes observed in the connectome are temporally linked to the activation of the LC-NE system and occur within minutes after intravenous injection of clozapine in mice expressing hM3Dq.

### Increased functional connectivity is dependent on LC-NE signaling

We next tested whether the DREADD-induced connectivity changes could be pharmacologically blocked by medetomidine, a selective agonist of the inhibitory adrenergic *alpha-2* receptor. Through *alpha-2* autoreceptors, medetomidine suppresses LC firing and NE release^42,43^. After pre-treating mice with a bolus injection of medetomidine (0.05 mg/kg, i.v.), medetomidine was also continuously infused at 0.1 mg/kg/h i.v. to maintain its levels stable throughout the ensuing rs-fMRI session, as previously described^44^. After 15 minutes of baseline recording, clozapine was again administered as described above (Figure 2A). Across all edges, Cohen’s D did not vary significantly over-time, showing that medetomidine prevented DREADD-induced LC activation (Figure 2D, E).

To examine the effect of different anesthesia regimes on FC over-time, we compared the effect size in mCh mice between both anesthesia conditions (i.e. 1% isoflurane vs. 0.5% isoflurane+medetomidine). We found no significant differences between the two experimental conditions in the baseline and transient periods (**Figure S2**). However, Cohen’s D was slightly increased in the active period, suggesting a reduction of connectivity across all edges in the isoflurane group. This is a well-known effect, most likely caused by the accumulation of isoflurane over-time^45^. The net effect was, however, null-to-small (average: +0.04, min: −0.04, max: +0.129) and about 7 times smaller than the DREADD-driven effect. Collectively, our findings indicate that the substantial changes in connectivity after DREADD activation causally depend on activation of the LC-NE system, as they are induced by LC-specific stimulation, and blocked by an *alpha*-2 adrenergic agonist.

### Spatial reconfiguration of the functional connectome after LC activation reflects transcript levels of adrenoceptors

Next, we mapped the location of the connections that were altered following LC activation. To this end, we compared the connectome matrix obtained from the first 15 minutes of the scan (baseline period) to the last 15 minutes of the scan (post clozapine, *‘active* period’). The analysis of the functional connectome after LC-NE activation revealed a large group of edges that display increased FC in the hM3Dq-mCh animals compared to mCh after clozapine injection (218 edges, time × group interaction, non-parametric, randomized permutation testing, family-wise error (FWE) corrected with Network Based Statistics at *p*<0.05) (Figure 3A). The spatial distribution of these hyper-connected edges is widespread and involves 64% of the 165 brain regions considered in the analysis (105 ROIs) (Figure 3B), which is in line with the widespread afferent fibers originating from the LC^46,47^.

**Figure 3.**
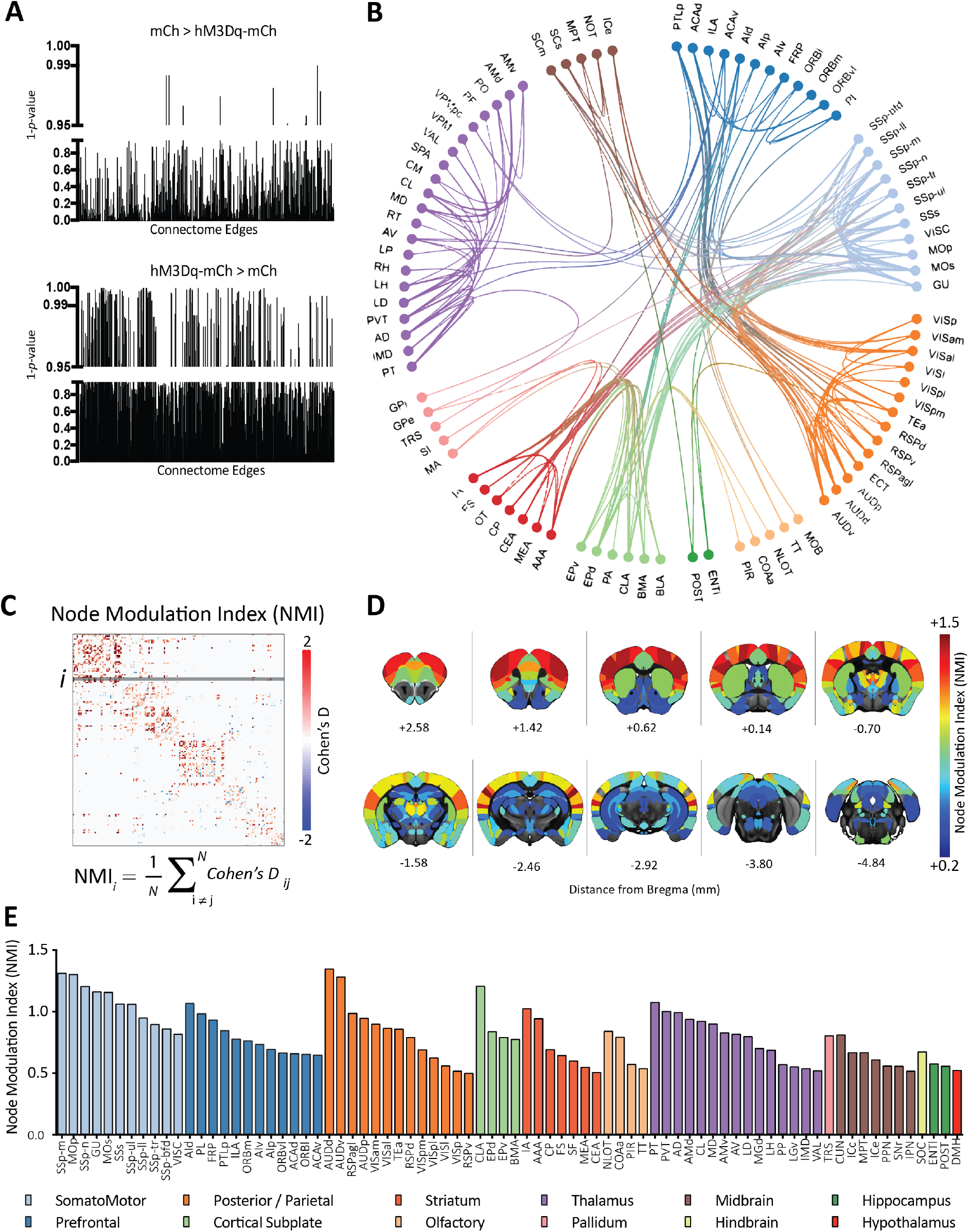
A whole-brain map showing the *connectome* after DREADD-induced LC activation. (**A**) Randomized non-parametric statistics report a drastic shift toward hyper-connectivity in the hM3D(Gq)-mCh group after clozapine injection (15 min bins). (**B**) Circos-plot showing the anatomical location of hyper-connected edges in response to LC activation (*p*<0.05, Network Based Statistics, FWE corrected). (**C**) Node Modulation Index (NMI), calculated as the averaged effect-size in each brain area (165 ROIs, based on the Allen Common Coordinate Framework). (**D**) Rendering of NMI in Allen MRI space, revealing a heterogeneous distribution across brain regions. (**E**) Bar plots representing all the ROIs with an NMI > 0.5 (moderate-to-strong effect). Full ROI list in Supplementary Table 1.

We quantitatively ranked the brain regions based on how LC activation changed the strength of its connections. To do this, we derived a surrogate ‘node modulation index’ (NMI) by measuring the averaged effect size for any brain region in the connectome matrix (Figure 3C). Note that only the top-10% of the connectome edges, based on FC strength at baseline, were considered in the analysis (see **Figure S3A**-C for a more detailed evaluation of other sparsity levels). The NMI varied across the brain (Figure 3D, E, full list in supplementary table 1). The strongest variations occurred in regions that are densely innervated by the LC, such as the primary sensory and somatomotor areas^48^, the claustrum^49^, the prefrontal cortex (notably the agranular insular cortex; pre- and infralimbic cortices; the frontal pole; and anterior cingulate areas)^16,26^, several nuclei of the amygdala^15^, the thalamus^19^, and the association cortex^26^ (Figure 3D, E).

The LC is a bilateral structure, and each locus primarily projects ipsilateral. Thus, we tested whether our LC stimulation would similarly affect the spatial patterns of FC within the left and the right hemispheres. We found remarkably similar changes in FC (**Figure S4**) on each side. The distribution of the effect size for all edges is right-skewed in both left and right hemispheres (**Figure S4A, B**), demonstrating increased connectivity after LC activation. Further, the changes in each node expressed with the NMI were highly linearly correlated between left and right hemisphere (Pearson’s correlation = 0.7764, *p*<0.0001, **Figure S4C**).

We then took advantage of the unique availability of molecular data in the same mouse strain, and asked if the anatomical heterogeneity found in the connectivity-based NMI maps relates to the spatial distribution of adrenergic receptors in the mouse brain. Gene transcript maps of adrenoceptors *alpha-1* (subunits a-d), *alpha-2* (subunits a-c), *beta-1* and *beta-2* were obtained from the Allen Mouse Brain Atlas^50^. The similarity with NMI maps was quantified using the Spearman rank correlation coefficient *(Rho)* across all 165 brain areas of the Allen’s Common Coordinate Framework. After correcting for testing multiple independent hypotheses, we found that the transcriptional maps of all adrenoceptors display a significant correlation with the NMI, with the exception of *beta*-2 adrenergic receptors (Figure 4A-D). All these correlations display a higher than chance Spearman’s rho value (*p*<0.0001 against a null distribution generated using 10.000 random permutations). We additionally compared our MNI maps against all dopamine (D1, D2, D3, D4, D5) and serotonin (Htr1, Htr2, Htr3, Htr4, Htr5, Htr6, Htr7) receptors, to assess the specificity of our results. We found a significant Spearman’s correlation with both D1 (*ρ*=0.4014, p_FDR_=118e^−7^) and D4 (*ρ*=0.3805, p_FDR_=137e^−6^) receptors, which survived permutation testing and FDR correction, but not with D2, D3 and D5 (**Figure S5**). None of the serotonin receptors had correlations higher than chance (**Figure S6**). These results represent the first brain-wide FC mapping in response to LC activation, revealing an anatomically specific *connectomic fingerprint* of LC hyperactivity, which maps well onto the spatial distribution of specific adrenoceptors and dopamine receptors.

**Figure 4.**
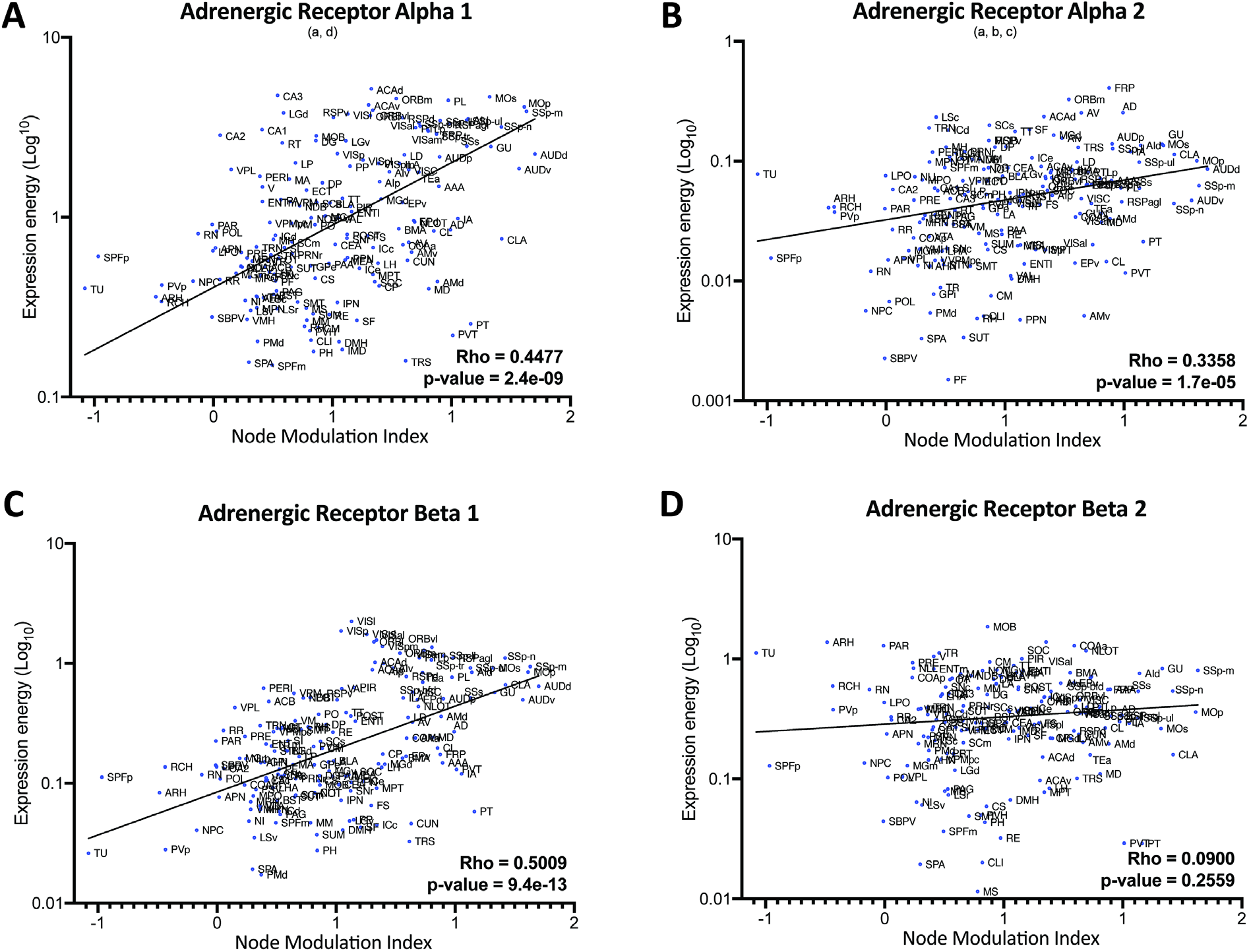
Functional connectivity changes after LC-NE activation spatially correlate with adrenoceptor gene expression. Spearman correlation coefficients, Rho, and associated p-value (FDR corrected) between Node Modulation Index and the transcriptional maps of genes coding (**A**) adrenoceptors alpha 1; (**B**) adrenoceptors alpha 2; (**C**) adrenoceptors beta 1; (**D**) adrenoceptors beta 2. Gene expression data was obtained from the Allen Mouse Brain Atlas (AMBA) and measured using *in situ* hybridization. Transcriptional levels across a macroscopic cortical area were summarized and plotted as the mean ISH intensity across voxels of that brain area, or ‘expression energy’.

### LC increased connectivity within large-scale resting-state networks

Thus far, our data driven analyses were focused on changes in connectivity between anatomically-defined pairs of brain regions. Next, we examined whether LC activation prompts large-scale resting-state network (RSN) reconfiguration, by calculating the spatial extent and connectivity strength in thirteen maximally-independent RSNs. These RSNs were obtained from an independent cohort of wild-type mice using a data-driven probabilistic independent component analysis (ICA), followed by an estimation of network coupling strength at the voxel level using a dual regression approach^51^ (for a complete list and spatial distribution of the networks please refer to our previous study^52^). Statistical analyses were conducted both at the voxel level (using non-parametric permutation testing with Threshold-Free Cluster Enhancement to correct for multiple comparisons which allows voxel-wise inference), and by measuring a connectivity-strength index within each RSNs (using a Linear Mixed-Model, see^52^).

In 6 out of 13 RSNs, we found significant group x time interactions resulting from an increase of within-network connectivity in the hM3Dq-mCh group, which was in stark contrast to the slight decrease in connectivity observed in the mCh group (Figure 5). The latter effect is likely attributable to an accumulation of isoflurane over time^45^ (see **Figure S2**) which is, however, clearly reversed upon LC activation. We observe the strongest differences in two networks previously linked to LC activity in humans exposed to stress^29^: (1) the Salience Network (hyperconnectivity in agranular insular area, anterior cingulate, ventro-medial striatum, accumbens, globus pallidus, parafascicular nucleus of the thalamus and hippocampus; Figure 5A), and (2) the Amygdala Network (hyperconnectivity in basomedial and basolateral amygdala, claustrum, sub-thalamic nucleus and zona incerta; Figure 5B). Additionally, LC activation also increased connectivity within the Association/Auditory Network (Figure 5C), the Dorsal Hippocampus Network (Figure 5D), the Striato-Motor Network (Figure 5E), and the Antero-Posterior Retrosplenial / Default-Mode-Network (Figure 5F). These results suggest that LC activation expands the synchrony of signals in several large-scale networks, most significantly in the Salience and Amygdala Networks. These results are remarkably similar to those in humans, showing that acute stress increases FC within homologous salience/amygdala networks^30,31^ in a beta-adrenergic receptor-dependent manner^31^.

**Figure 5.**
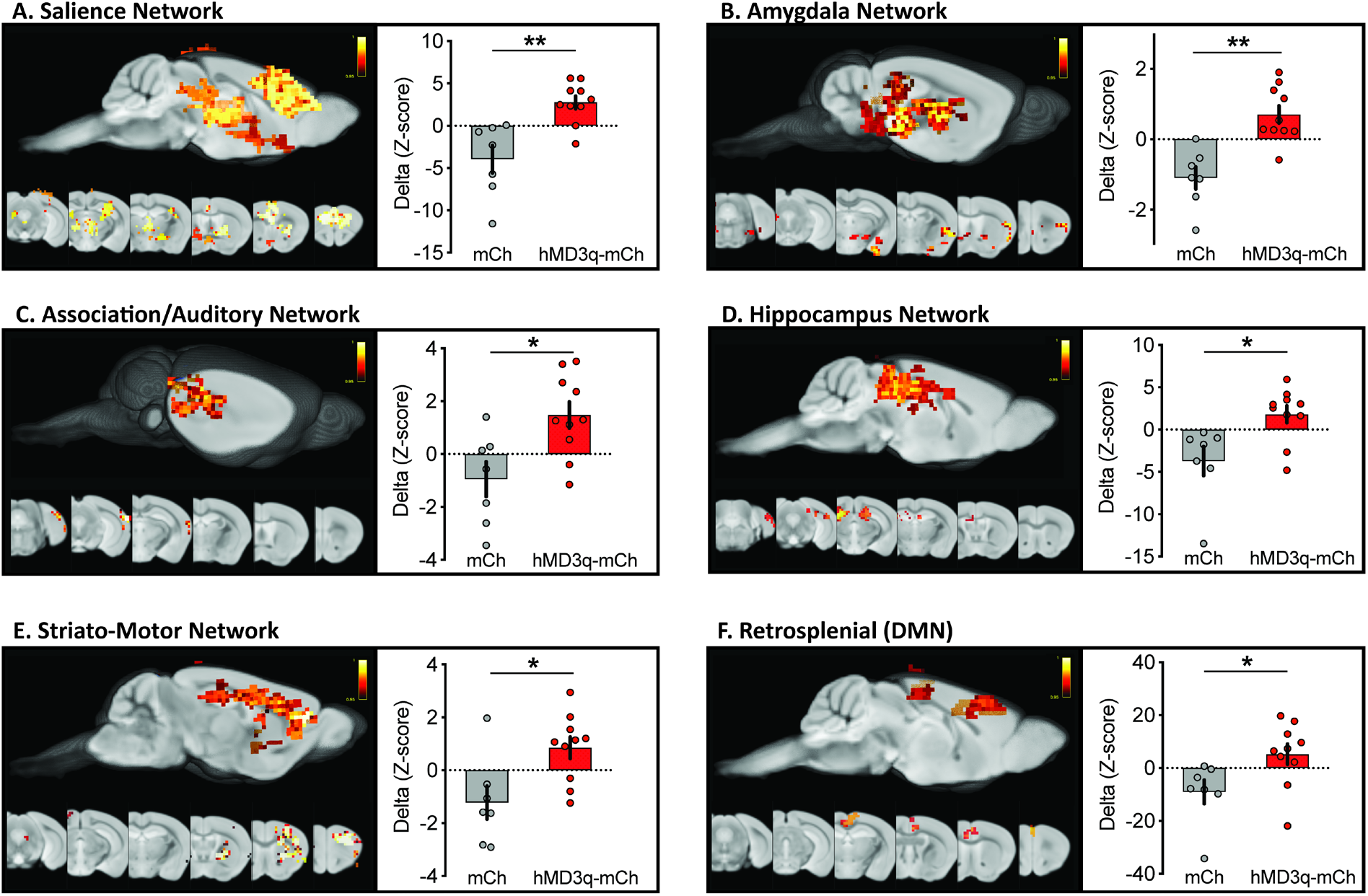
Rapid changes in resting-state network connectivity after LC activation. Voxel-wise dual-regression analysis of revealed clusters of significant ‘group x time interactions’ in 6/13 resting-state networks (*p*-value<0.05, TFCE corrected). A linear mixed model showed significant network strength increases in the hM3Dq-mCh group compared to mCh control mice, after clozapine injection. Bar plots represent mean ± SEM. *=*p*<0.05, **=*p*<0.01 (FDR corrected)

### Network connectivity changes correlate with norepinephrine and dopamine turnover

After completion of the two rs-fMRI sessions, mice were given one/two weeks to recover (see methods). Eight hM3Dq-mCh mice and five mCh controls were injected again with clozapine (0.03 mg/kg, i.p.), and their brains were collected 90 minutes later to address two important issues; first, we wanted to ensure that clozapine exclusively activated noradrenergic LC neurons, and that activation occurred in all hM3Dq-mCh mice but not in the mCh controls. Second, we wanted to assess whether LC activation led to measurable NE release in target regions throughout the brain, and whether NE levels would correlate with observed changes in network connectivity. In order to address these two issues in each individual mouse, we pursued a two-pronged strategy for tissue processing (Figure 6A): Freshly collected brains were split with a razorblade along the superior colliculus to collect one section containing the LC for immunohistochemistry, and a second section containing the forebrain. From the forebrain section, we rapidly dissected cerebral cortex, hippocampus and dorsal striatum on ice. Samples were snap frozen and processed for analysis of monoamines and their metabolites using reversed-phase ultra-high-performance liquid chromatography (uHPLC) coupled with electrochemical detection (Figure 6A).

**Figure 6.**
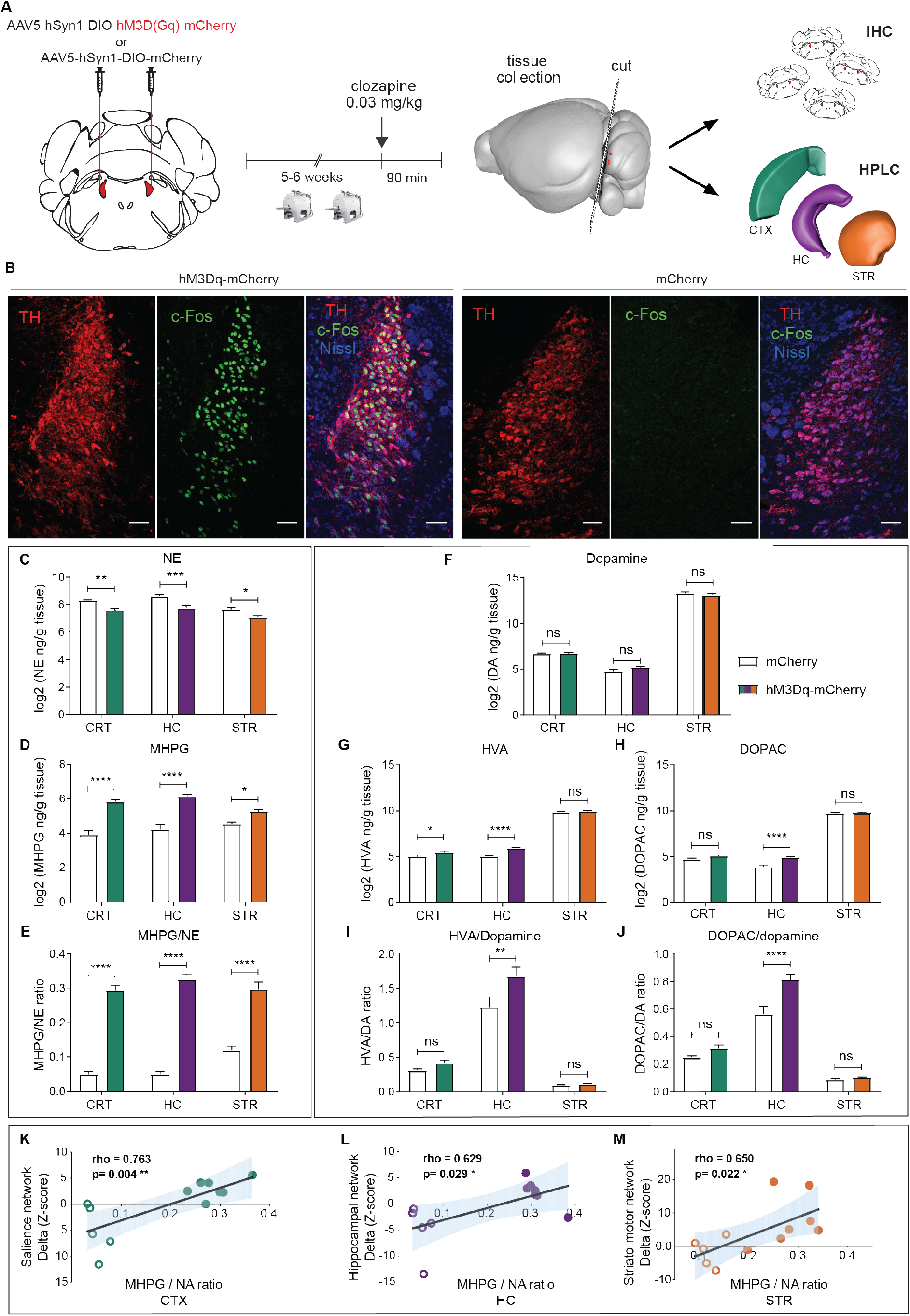
NE and DA turnover induced by LC activation correlates with rs-fMRI data. (**A**) Outline of the tissue collection strategy following completion of fMRI scans. (CTX: cortex, HC: hippocampus, STR: striatum = caudate putamen) (**B**) Representative images of cFos expression in the LC after clozapine injection in hM3Dq-mCh or mCh mice. All scale bars: 50 μm. (**C-J**) Levels of monoaminergic neurotransmitters and their metabolites in the cortex, hippocampus and striatum. NE was reduced in all brain regions of hM3Dq-mCh mice relative to mCh controls (C: main effect of group: F(1,33)=42.48, *p*<0.0001), MHPG was increased (D: main effect of group F(1,33)=121.80, *p*<0.0001) and NE turnover ratio (MHPG/NE) was increased (E: main effect of group: F(1,33)=291.50, *p*<0.0001; two-way ANOVA with Sidak *post hoc* tests). Although there was no difference in the levels of DA (F: main effect for group F(1,33)=0.81, *p*=0.3748), its metabolite HVA was increased in the cortex and the hippocampus in hM3Dq-mCh mice (G: significant main effect for group F(1,11)=11.60, *p*=0.0059) and DOPAC was increased only in the hippocampus (H: significant main effect for group F(1,33)=22.13, *p*<0.0001;). The DA turnover ratios were only increased in the hippocampus, for both HVA/DA (I: main effect of group: F(1,33)=8.91, *p*=0.0053), and DOPAC/DA (J: main effect of group: F(1,33)=19.81, *p*<0.0001), two-way ANOVA with Sidak *post hoc* tests). (**K-M**) Correlation between FC changes and neurotransmitter levels in the cortex (**K**), in the hippocampus (**L**), and in the striatum (**M**). Circles correspond to mCh mice, filled circles to hM3Dq-mCh mice. *p<0.05, **p<0.01, ***p<0.001, ****p<0.0001. Data represent mean ± SEM.

Co-labelling for TH and cFos revealed that clozapine injection only induced a strong and reliable activation of noradrenergic neurons in the LC of the hM3Dq-mCh mice (representative image in Figure 6B, sections of all mice presented in **Figure S7**). This validates that LC activation was successful in the mice undergoing fMRI scans. In parallel, we used uHPLC to measure and quantify the monoaminergic neurotransmitters norepinephrine (NE), dopamine (DA) and serotonin (5HT), as well as their main metabolites 3-methoxy-4-hydroxyphenylglycol (MHPG, metabolite of NE), homovanillic acid (HVA, metabolite of DA), 3,4-dihydroxyphenylacetic acid (DOPAC, metabolite of DA), and 5-hydroxyindoleacetic acid (5-HIAA, metabolite of 5HT). We were able to reliably detect and quantify all measured compounds (see representative chromatographs in **Figure S8**). NE levels decreased in all brain regions (Figure 6C), suggesting that LC activation for 90 minutes had depleted NE storage vesicles, as would be expected after sustained high-frequency firing. In agreement, MHPG levels increased in all brain regions (Figure 6D), resulting in a very strong increase in catabolic NE turnover (MHPG/NE, Figure 6E). Dopamine levels were not changed in any of the brain regions under investigation (Figure 6F), yet we observed a robust increase in both hippocampal HVA and DOPAC (Figure 6G, H), and a small but significant increase of HVA in the cortex (Figure 6G). Dopamine turnover ratios (HVA/DA and DOPAC/DA) were increased in the hippocampus but not in striatum or cortex (Figure 6I, J). These results are in line with recent evidence that LC neurons can release DA in certain brain regions including the hippocampus^10,11,19,53^. Our data newly suggest that DA release from LC neurons may be biased towards the hippocampus compared to the cortex and striatum. Epinephrine, 5-HT and 5-HT turnover ratios were not altered in any of the brain regions sampled (**Figure S9**).

As we performed fMRI and uHPLC analyses in the same mice (although with a temporal delay of 1 week and in response to separate injections with clozapine), we were able to conduct a correlation analysis between individual differences in neurotransmitter turnover and corresponding changes in network connectivity. We found positive correlations between the NE and DA turnover ratios in the cortex and changes in FC within the Salience Network (Figure 6K, **Figure S10A**), as well as between NE and DA turnover ratios in the hippocampus and FC in the Hippocampus Network (Figure 6L, **Figure S10E**). NE turnover in the striatum, but not DA turnover, correlated with FC changes in the Striato-Motor Network (Figure 6M, **Figure S10C, F**). Importantly, we observed no correlation between 5-HT levels/turnover and the respective network connectivity changes in any of the brain regions tested (**Figure S10G-I**). These results collectively suggest that brain network changes we observe with rs-fMRI are tied to the amount of NE and/or DA released in a given region.

### LC neurons project sparsely to the dorsal striatum (caudate-putamen)

Although increased NE turnover in the cortex and hippocampus was expected due to strong innervation by the LC, the increased NE turnover in the dorsal striatum (consisting of caudate and putamen)^54^ was surprising, because this region is widely thought to be devoid of NE projections^24,46^. In agreement with increased striatal NE turnover, we also observed a strong increase in the node modulation index (NMI) in the caudate-putamen (Figure 3E), and increased functional connectivity within the large-scale Striato-Motor Network (Figure 5E). Thus, we decided to first test whether we could detect noradrenergic axons within the caudate-putamen. We stained for the norepinephrine reuptake transporter Slc6a2 (NET), which is expressed exclusively in noradrenergic cells^55,56^. Although the caudate-putamen appears devoid of NET compared to the intense NET staining seen in the adjacent cortex (Figure 7A), we clearly detected long, thin axons in the caudate-putamen (Figure 7B). To investigate whether these axons originate from LC neurons, we delivered a retrograde AAV2 virus^57^ carrying floxed mCherry into the dorsolateral caudate-putamen of DBH-iCre mice. Seven weeks after virus injection, we stained the LC and detected a clearly recognizable subset of LC neurons that expressed mCherry in transgenic animals (n=3) but not in wild-type controls (n=1) (representative images from each animal shown in Figure 7C and **Figure S11**). We were able to reproduce these findings with a second Cre-dependent retrograde AAV2 virus that expresses EGFP (Figure 7D), which also allowed us to detect axons in the caudate-putamen that were co-labelled with both EGFP and NET (Figure 7E). Together, these data show that there are sparse projections from LC to the caudate-putamen. These projections could account for the increase in NE levels detected in the striatum after LC stimulation, as well as for the increased functional connectivity observed in the caudate-putamen and in the large-scale Striato-Motor Network.

**Figure 7.**
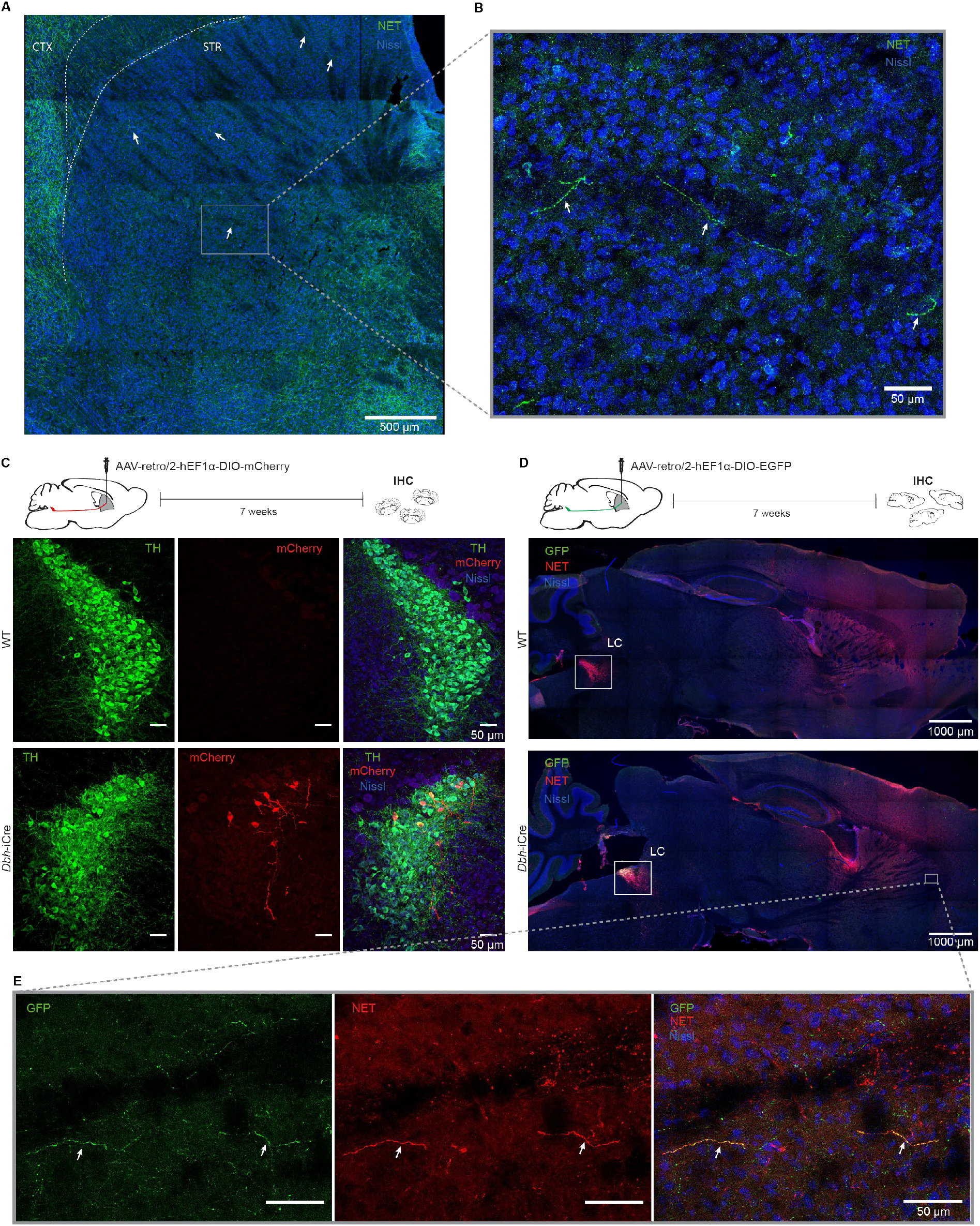
The dorsal striatum (caudate-putamen) is innervated by the LC. (**A**) Immunohistochemical localization of the norepinephrine transporter (NET) reveals noradrenergic axons in the caudate-putamen (**arrows**). (**B**) Magnification of the box in (**A**) showing NET+ axons (arrows). (**C**) A retro-AAV2 that expresses Cre-dependent mCherry was delivered to the dorsolateral caudate-putamen and resulted in mCherry+ neurons in the LC of DBH-iCre (n=3), but not wild type (n=1) mice (representative pictures from one animal shown, additional images for all mice in **Figure S11**). (**D**) Stereotactic delivery of a Cre-dependent retro-AAV2 virus expressing EGFP in the dorsolateral caudate-putamen resulted in EGFP+ LC neurons in a DBH-iCre mouse, but not in a wild type mouse. (**E**) Magnification of the box in (**D**) showing the axonal co-localization of NET and EGFP in the caudate putamen of a DBH-iCre animal. CTX=cortex; STR=striatum

## DISCUSSION

### Revealing the connectomic fingerprint of norepinephrine release after LC activation

Neuromodulatory systems of the brain track and integrate environmental signals, exert powerful control over neuronal function, and are the primary targets of most neuropsychiatric treatment strategies^58–60^. It is, however, challenging to study the impact of an individual neuromodulatory system across large-scale neuronal networks. The advent of optogenetics and chemogenetics, combined with advances in rodent imaging capabilities, now enables us to link circuit-level manipulations to global network changes. Recent studies have used these approaches to show that selective manipulation of dopamine^61,62^ and serotonin^63^ release can lead to global activity changes measured by fMRI. Our work extends these studies by assessing the role of the LC-NE system in assembling and rearranging connectivity within and between well-defined large-scale neuronal systems using rs-fMRI. This task-free, system level approach allows interrogation of different functional circuits in parallel (i.e. the *connectome*), and is therefore well-suited for testing the concept that changes in neuromodulator levels can shift neural ‘brain states’. Further, rs-fMRI is emerging as a tool for translational neuroscience, as it can be performed in multiple species: humans^22^, primates^64^, and rodents^65^. However, clinical studies using rs-fMRI often do not translate well into useful diagnostic or prognostic information for individual patients^66^, in part because it is unclear how molecular and/or cellular defects are reflected by changes in connectivity at the network level. Our novel chemo-connectomic approach attempts to bridge this gap by leveraging the full power of genetic and molecular tools available in mice. Additionally, it establishes the framework necessary to link the activity of neuromodulatory systems to clinically relevant brain signals and their neuroanatomical substrates.

### Adrenoceptor distribution as an organizing principle affording “global specificity”

Neurons in the LC are topographically organized based on their widespread efferent projections^46,47^, and circuit-based approaches reveal functional specificity of sub-populations of LC neurons based on their target projection regions^5,12–16,18,19,67,68^. LC neurons appear to form neuronal ensembles, which can be activated in response to isolated sensory stimuli^67,69^. With increasing stimulus strength, more LC-ensembles can be activated, leading to “global” LC activation in response to strong stimuli (such as stress exposure)^69^. Widespread NE release in response to this global LC activation is commonly thought to act as a broadcast signal, which modulates brain states, gearing specific networks towards integrating environmental information, allowing to select/trigger adequate behaviors^1,12,47^. However, it remains unclear how such a global signal can achieve network-wide specificity (e.g. to interrupt ongoing behavior and trigger an orienting response). The widespread projections of the LC make it unlikely that the anatomical connections of these afferents encode this specificity^47^. In line with this assumption, a study using manganese-enhanced MRI suggested that the anatomy of LC projections alone does not predict functional connectivity^70^. However, NE release in projection regions can be mediated by adrenoceptor distribution. This was, for example, shown in the basal forebrain, where *alpha*-1 and *beta* −1 adrenoceptors are expressed on cholinergic neurons, whereas *alpha*-2 receptors reside on GABAergic interneurons. Thus, NE release can simultaneously activate cholinergic neurons and inhibit GABAergic neurons^71,72^. To address the potential modulatory role of adrenoceptors in the effects of global LC activation, we correlated brain-wide changes in functional connectivity (FC) with gene expression density of adrenergic, dopaminergic and serotonergic receptors. We observed strong correlations between FC and expression levels of *alpha*-1 and *beta*-1 adrenoceptors, which are known to have low binding affinity for NE and are thus only activated in response to high levels of NE release^24,26^. A weaker correlation was also found for inhibitory *alpha*-2 receptors. The absence of a correlation with *beta*-2 adrenergic receptors was unexpected, given that both *beta*-1 and *beta*-2 adrenoceptors are ubiquitously expressed and have similar binding affinity for NE^73^, although contrasting effects of *beta*-1 and *beta*-2 receptors on working memory performance have previously been reported^74,75^. To the best of our knowledge, our results are the first to demonstrate a role for adrenergic receptor distribution in providing specificity to global NE release on a brain-wide, functional level. This helps to explain how a “global broadcast” signal can achieve highly specific functional and behavioral effects. Molecular neuroscience tools in mice are rapidly advancing, for example recent work has conducted careful input-output tracing of LC connectivity using novel trans-synaptic tracing tools^76^, and refined sequencing approaches have begun to genetically track LC projections across the mouse cortex^77^. As these and similar datasets become catalogued and readily accessible, our rs-fMRI data can serve as a resource to map functional, brain-wide connectome data onto the molecular characteristics of the LC-NE system.

### LC activation recapitulates many of the complex effects triggered by stress exposure

Modern theories of LC function propose that LC activation serves to optimize the trade-off between exploitation and exploration, with strong, global LC activity causing interruption of ongoing activity to enable the selection of appropriate behaviors^20^. Global LC activity is robustly triggered by noxious/stressful stimuli^12,13,24,25,67,78^, which induce anxiety and reduce exploratory activity, through circuits involving the amygdala and prefrontal cortex^3,13–16^. Our DREADD-induced activation of the LC similarly reduces exploratory activity and increases anxiety, and globally induces cFos expression throughout the LC. Therefore, our global LC activation likely resembles peak LC activity that would normally be triggered by stressful stimuli.

Our data are well in line with findings in humans where acute stress exposures (induced by aversive movies or social stressors) were followed by a significant increase of FC in the Salience Network^30^ and Default Mode Network^79^. Similarly, task-based fMRI reveals that acute stress leads to increased interconnectivity and positive BOLD responses within several cortical regions related to salience processing (frontoinsular, anterior cingulate, inferotemporal, and temporoparietal) and subcortical regions (amygdala, striatum, thalamus, hypothalamus, hippocampus and midbrain) as a function of stress response magnitude^31,80–82^. Notably, increased connectivity in the Salience Network was blocked by systemic administration of a *beta-adrenergic* receptor antagonist (propranolol)^31^. Given that LC activation is only one aspect of the highly complex changes observed during an acute stress response^83^, and considering that our analyses were performed in lightly anesthetized mice, it is remarkable that our results for the Salience Network, the Amygdala Network, and also the Default-Mode Network closely resemble fMRI and rs-fMRI findings described after acute stress exposures in humans^29,82^ (Figure 5). Since our model allowed us to selectively and directly manipulate LC activity with DREADDs, our results causally show that selective LC activation rapidly reorganizes functional connectivity within specific large-scale networks.

Long-lasting hyperactivity of the LC is often observed after severe or chronic stress exposures^78,84^ and is considered a hallmark feature of post-traumatic stress disorder (PTSD)^4,85^. Two recent rs-fMRI studies showed - in both rats and mice - that chronic stress exposure resulted in large-range increases in functional network connectivity for regions including prelimbic/infralimbic areas, amygdala, cingulate cortex and hippocampus^86,87^. In PTSD patients, fMRI reveals network wide changes in the amygdala, insula, hippocampus and anterior cingulate cortex^88,89^. Therefore, the changes observed after chronic stress exposure in rodents and in PTSD patients are strikingly similar to the effects of selective LC stimulation. Hyperactivity of the LC represents only one aspect of the multifaceted changes occurring in stress-related pathologies, but our data provide additional evidence that modulating LC activity might be a promising therapeutic approach^78,90^.

### Functional connectivity after LC activation involves dopamine release and dopamine receptors

Several recent reports have shown that dopamine (DA) can be co-released from LC terminals in the hippocampus^10,11,53^ and thalamus^19^. We also detect a strong increase in DA turnover in the hippocampus, and a subtle increase in the cortex (with no effects in the striatum). Our cortex samples contained heterogeneous regions, it is likely that specific cortical subregions (e.g. mPFC) display stronger changes in DA turnover, similar to the effects observed in the hippocampus. In concordance with our data, previous microdialysis experiments have detected release of both DA and NE in the medial prefrontal cortex after chemical or electrical activation of the LC, while the same stimulations increased NE but not DA in the nucleus accumbens and caudate nucleus^91–93^. In agreement with this regional specificity of NE and DA release, our study reveals positive correlations between cortical DA turnover and FC within the Salience Network, as well as between hippocampal DA turnover and FC within the Hippocampus Network. In contrast, no significant correlations were detected in the striatum. Furthermore, brain-wide changes in FC induced by LC stimulation correlated positively with the transcript expression levels of D1, D3 and D4 dopamine receptors. Together, these data provide new evidence for a physiologically relevant, regionally restricted role of DA release in response to LC stimulation. Future studies will have to test whether all LC neurons (or only selected sub-populations with specific projection targets) are able to release DA, and if specific interactions at the projection site are required to enable DA release. Finally, it remains unclear whether this co-release occurs under physiological conditions, as evidence suggests that it depends on the firing rate of individual LC neurons^93^.

### LC-induced synchronization in the dorsal striatum (caudate-putamen)

Although close interactions between dopaminergic and noradrenergic systems have long been recognized^94^, the dorsal striatum (caudate-putamen) is widely thought to be devoid of noradrenergic projections^24,46^, and indeed staining for the NE transporter (NET, Figure 7A) or DBH^95^ shows a strong depletion in the caudate-putamen. Given these data, we were surprised when our connectome analysis revealed that LC activation strongly increased the strength of connectivity of the caudate-putamen (Figure 3C). Further, we observed synchronization of rs-fMRI activity in the Striato-Motor Network, which positively correlated with increased NE turnover in the same region (Figure 6M). Indeed, several studies have shown fairly high levels of NE^96^ in the caudate-putamen, as well as extracellular striatal NE increase after mild stress (handling)^97^ or LC stimulation^92^. Adrenergic *alpha*-1, *alpha*-2 and *beta*-1 receptors are abundantly expressed on striatal pre- and post-synaptic membranes and cell bodies^98–102^, and *beta*-adrenergic receptor binding density is very high in the caudate-putamen of rodents and humans^103–105^. Our results show that a considerable number of LC neurons directly project to the caudate-putamen, which is in line with early retrograde tracing work^106^ but was rarely recognized in more recent literature. Taken together, the fact that the caudate-putamen expresses high levels of adrenoceptors, is sparsely innervated by LC neurons, and contains increased NE levels after LC stimulation, suggests that the strong increase of functional connectivity in the caudate-putamen after LC stimulation is driven by direct projections from the LC. These findings may be relevant for Parkinson’s disease, where an involvement of the LC-NE system is increasingly being recognized^8,107^.

### Conclusions

Using the novel chemo-connectomics approach presented here, we provide the first brain-wide analysis of connectome reconfiguration in response to selective LC activation. We show a profound, rapid and specific activation of large-scale networks related to salience processing, which occurs within minutes after clozapine administration specifically in mice expressing DREADDs. These effects are observed similarly in each hemisphere and they are blocked by activating presynaptic, auto-inhibitory *alpha-2* adrenoceptors. Shifts in large-scale network connectivity correlate spatially with the distribution of adrenergic and dopaminergic receptor levels and with (post-mortem) measurements of NE, DA, and their metabolites in a within-subject design. These network effects are accompanied by increased pupil size and heightened anxiety, suggesting that the observed changes in brain network organization ultimately serve to promote vigilance and threat detection.

## Acknowledgements

We thank Han-Yu Lin for genotyping, help with image analysis and lab management and the team of the EPIC animal facility for providing animal care. We thank Jean-Charles Paterna from the Viral Vector Facility (VVF) of the Neuroscience Center Zurich, a joint competence center of ETH Zurich and University of Zurich for producing viral vectors and viral vector plasmids. We thank Prof. Isabelle Mansuy for generously providing support and space in her lab during the early stages of this project. We thank Dr. Ben Fulcher for providing the gene-expression data and help with the correlational analyses, and Pierre-Luc Germain for feedback on statistical analyses. We thank Tilo Aurelio Gschwind for providing guidance for cFos immunohistochemistry, and Ladina Hösli for providing helpful advice during the Rotarod experiments.

## Methods

### Mice

C57Bl/6J male mice (2 months old) were obtained from Janvier (France). Heterozygous C57BL/6-Tg(Dbh-icre)1Gsc mice^33^ were generously provided by Prof. Günther Schütz and kept in breeding trios with wild-type C57Bl/6J mice at the ETH Zurich animal facility (EPIC). Mice were maintained in IVC cages with food and water ad libitum, in a temperature- and humidity-controlled facility on a 12-hour reversed light-dark cycle (lights off: 8:15am; lights on: 8:15pm). The experimenters were blinded to the DREADD groups for all experiments. All experiments were performed in accordance with the Swiss federal guidelines for the use of animals in research, and under licensing from the Zürich Cantonal veterinary office.

### Outlier Removal

All mice (hM3Dq-mCh n=11; mCh n=8) successfully completed both fMRI sessions. However, due to a fault in the ventilation system that artificially supports breathing after the MRI, three animals died after the second session (hM3Dq-mCh n=2; mCh n=1). For all the others, the time until conscious intention to move was 10.6±4 minutes. One animal (mCh group) was excluded from the rs-fMRI analysis because the preparation time exceeded 30 minutes (average time between anesthesia induction and start of the MRI scanning, including intubation and cannulation was 12.8±3 minutes). In the weeks between the second fMRI session and tissue collection for molecular analyses, 2 mice were found dead in their cage (n=1 from hM3Dq-mCh, and n=1 from mCh groups). Otherwise, no mice were excluded from any of the experiments.

### Stereotaxic brain injections

Viral vectors and viral vector plasmids were designed and produced by the Viral Vector Facility (VVF) of the Neuroscience Center Zurich. The viruses used had a physical titer of 6.0-6.5 x 10^12^ vg/ml. For virus delivery, 2 to 3 month old mice were subjected to stereotactic brain injections. The mice were anesthetized with isoflurane and placed in a stereotaxic frame. For analgesia, animals received a subcutaneous injection of 2mg/kg Meloxicam and a local anesthetic (Emla cream; 5% lidocaine, 5% prilocaine) before and after surgery. A pneumatic injector (Narishige, IM-11-2) and calibrated microcapillaries (Sigma-Aldrich, P0549) were used to inject 1 μL of virus (either ssAAV-5/2-hSyn1-dlox-hM3D(Gq)_mCherry(rev)-dlox-WPRE-hGHp(A) or ssAAV-5/2-hSyn1-dlox-mCherry(rev)-dlox-WPRE-hGHp(A)) bilaterally into the locus coeruleus (coordinates from bregma: anterior/posterior-5.4 mm, medial/lateral +/-1.0 mm, dorsal/ventral-3.8 mm). For the retrograde AAV2 injection, 0.8 μL of ssAAV-retro/2-hEF1α-dlox-hChR2(H134R)_mCherry(rev)-dlox-WPRE-hGHp(A) or ssAAV-retro/2-hEF1a-dlox-EGFP(rev)-dlox-WPRE-bGHp(A) was delivered bilaterally to the dorsolateral site of the caudate-putamen (from bregma: anterior/posterior 0.86 mm, medial/lateral +/- 1.8 mm, dorsal/ventral −3.2 mm). The health of the animals was evaluated by post-operative checks over the course of 3 consecutive days. Brain illustrations were created with the Scalable Brain Atlas^50,108^.

### Pupillometry

For pupil recordings we used a Raspberry Pi NoIR Camera Module V2 night vision camera, an infrared light source (Pi Supply Bright Pi - Bright White and IR camera light for Raspberry Pi) and a Raspberry Pi 3 Model B (Raspberry Pi Foundation, UK). Animals were anesthetized with isoflurane (4% induction, 1.5% maintenance), then an intraperitoneal catheter delivering PBS was placed and the video recording was initiated. After 2 minutes of baseline recording, 0.03 mg/kg clozapine (Sigma-Aldrich, Steinheim, Germany) was injected through the catheter and the video recording continued for another 8 minutes. Pupil diameter was measured using a custom MATLAB (MathWorks, Natick, MA, USA) script. Frames were binarized and an ellipse-fitting algorithm was used to approximate the pupil size. Measurements after clozapine injection were then normalized to baseline (measurements before clozapine).

### Open field test (OFT)

Open-field testing took place inside sound insulated, ventilated multi-conditioning chambers (TSE Systems Ltd, Germany). The open field arena (45 cm (l) x 45 cm (w) x 40 cm (h)) consisted of four transparent Plexiglas walls and a light grey PVC floor. For all experiments, mice were tested under dim lighting (4 Lux across the floor of the open field, provided by four equally spaced yellow overhead lights) with 75-77 dB of white noise playing through the speakers of each box, as described previously^39^. The room housing the multi-conditioning chambers was illuminated with red LED lights (637 nm). Animals were injected i.p. with 0.03 mg/kg clozapine (Sigma-Aldrich, Steinheim, Germany) and placed directly into the center of the open field. Tracking/recording was initiated upon first locomotion grid beam break, lasted for 30 minutes and was analyzed in 3-minute bins.

### Tissue preparation, antibodies and immunohistochemistry

For experiments where brain tissue was collected exclusively for immunohistochemistry, mice were deeply anesthetized with pentobarbital (150 mg/kg, i.p.) and perfused intracardially through the left ventricle for 2 minutes, with approximately 20 mL ice-cold PBS (pH 7.4). The brain was dissected, blocked and fixed for 2-3 hrs in ice-cold paraformaldehyde solution (4% PFA in PBS, pH 7.4). The tissue was rinsed with PBS and stored in a sucrose solution (30% sucrose in PBS) at 4°C, overnight. Then the tissue was frozen in tissue mounting medium (Tissue-Tek O.C.T Compound, Sakura Finetek Europe B.V., Netherlands), and sectioned coronally using a cryostat (Leica CM3050 S, Leica Biosystems Nussloch GmbH) into 40 μm thick sections. The sections were immediately transferred into ice-cold PBS and stored at −20°C.

For immunohistochemistry, brain sections were submerged in primary antibody solution containing 0.2% Triton X-100, and 2% normal goat serum in PBS, and were incubated at 4°C under continuous agitation over 2 nights. Then the sections were washed 3 times in PBS for 10 minutes/wash, and transferred in secondary antibody solution containing 2% normal goat serum in PBS. After 3 more PBS washes, the sections were mounted onto glass slides (Menzel-Gläser SUPERFROST PLUS, Thermo Scientific), air-dried and coverslipped with Dako fluorescence mounting medium (Agilent Technologies). The primary antibodies used were: rabbit anti-mCherry (ab167453, Abcam, 1:1000), mouse anti-TH (22941, Immunostar, 1:1000), rabbit anti-cFos (226 003, Synaptic Systems, 1:5000), mouse anti-NET (NBP1-28665, Novus Biologicals, 1:1000), chicken anti-GFP (ab13970, Abcam. 1:1000). The secondary antibodies used were: goat anti-rabbit Alexa 546 (A11035, Life Technologies, 1:300), goat anti-mouse Alexa 488 (ab150113, Abcam, 1:300), goat anti-mouse Cy3 (115-165-003, Jackson ImmunoResearch, 1:300), goat anti-rabbit Alexa Fluor 488 (A-11008, Thermo Fisher Scientific, 1:500), goat anti-chicken (A-11039, Thermo Fischer Scientific, 1:1000) and Nissl stain (N21483, NeuroTrace 640/660 Nissl stain, Molecular probes).

Microscopy images were acquired in a confocal laser-scanning microscope (CLSM 880, Carl Zeiss AG, Germany), maintaining a pinhole aperture of 1.0 Airy Unit and image size 1024×1024 pixels. Images of LC were acquired using a Z-stack with a 20x objective and pixel size 0.59 μm. Images of the caudate putamen and the whole brain (sagittal plane) were acquired with the 20x (pixel size 0.59 μm) and the 10x (pixel size 1.19 μm) objective respectively, using a Z-stack and tiles. Images of axons were acquired with the 20x (pixel size 0.59 μm) or the 40x (pixel size 0.35 μm) objective.

### Tissue collection for uHPLC and immunohistochemistry

When brain tissue was collected for both uHPLC and immunohistochemistry, mice were rapidly euthanized by cervical dislocation. The brain was first divided into anterior and posterior with a single cut from a razorblade at the beginning of the cerebellum as shown in Figure 5A. The cortex (overlying the hippocampus), hippocampus and striatum were immediately dissected on an ice-cold glass surface, snap-frozen in liquid nitrogen and stored at −80°C until further processing for uHPLC. The posterior part including the locus coeruleus was fixed in 4% PFA for 2 hours, cryoprotected in a sucrose solution and frozen in mounting medium as described above for immunohistochemistry.

### MRI

Data acquisition was performed on a Biospec 70/16 small animal MR system (Bruker BioSpin MRI, Ettlingen, Germany) equipped with a cryogenic quadrature surface coil for signal detection (Bruker BioSpin AG, Fällanden, Switzerland). Standard adjustments included the calibration of the reference frequency power and the shim gradients using MapShim (Paravision v6.1). For anatomical assessment, a T2-weighted image is acquired (FLASH sequence, in-plane resolution of 0.05 × 0.02 mm, TE = 3.51, TR = 522ms). For functional connectivity acquisition, a standard gradient-echo echo planar imaging sequence (GE-EPI, repetition time TR=1s, echo time TE=15ms, in-plane resolution RES=0.22×0.2mm^2^, number of slice NS=20, slice thickness ST=0.4 mm, slice gap=0.1mm) was applied to acquire 2280 volumes in 38 min. After 15 minutes of GE-EPI acquisition, a bolus of 0.03 mg/kg clozapine was intravenously injected to activate DREADDs.

### Anaesthesia

The levels of anaesthesia and physiological parameters were monitored to obtain a reliable measurement of functional connectivity following established protocols^44,109^. Briefly, anaesthesia was induced with 4% isoflurane and the animals were endotracheally intubated and the tail vein cannulated. Mice were positioned on a MRI-compatible cradle, and artificially ventilated at 80 breaths per minute, 1:4 O_2_ to air ratio, and 1.8 ml/h flow (CWE, Ardmore, USA). A bolus injection of muscle relaxant (pancuronium bromide, 0.2 mg/kg) was administered, and isoflurane was reduced to 1%. Throughout the experiment, mice received a continuous infusion of pancuronium bromide 0.4 mg/kg/h. Body temperature was monitored using a rectal thermometer probe, and maintained at 36.5 °C ± 0.5 during the measurements. The preparation of the animals did not exceed 15 minutes. In an additional experiment, mice were pre-treated with a bolus injection of medetomidine 0.05 mg/kg, followed by a continuous infusion at 0.1 mg/kg/h.

### Resting-state fMRI data pre-processing and analysis

Resting state fMRI datasets were de-spiked and artefacts were removed using an existing automated pipeline, adapted for the mouse^109^. This procedure includes ICA-based artefact removal, motion correction and regression. Thereafter, data sets were band-pass filtered (0.01-0.25 Hz), skull-stripped and normalized to the Allen Brain Institute reference atlas (http://mouse.brain-map.org/static/atlas) using ANTs v2.1 (picsl.upenn.edu/ANTS).

Two types of analyses were performed on this data to compare the influence of LC activation on functional connectivity; first, we employed an exploratory and data-driven *connectome* analysis to describe the temporal and spatial changes at the whole-brain level. Briefly, BOLD time series are extracted using a subset of ROIs from the Allen’s Common Coordinate Framework (V3, http://help.brain-map.org/download/attachments/2818169/MouseCCF.pdf), which consisted of 165 ROIs from isocortex, hippocampal formation, cortical subplate, striatum, pallidum, thalamus, hypothalamus, hindbrain and midbrain (full list available in supplementary material, table 1). Connectivity couplings between all ROIs are measured using a regularized Pearson’s correlation coefficient implemented in FSLNets, using sliding time windows of either 1 minute (Figure 2) or 15 minutes (Figure 3). For each block, the resulting connectome matrices are fed into a nonparametric permutation testing with 5000 permutations to detect differences between groups, corrected for multiple comparisons with Network Based Statistics (NBS).

In our second analyses, we focused on changes in spatial patterns of correlated activity, also called *resting-state networks* (RSNs). We selected 15 meaningful RSNs from an independent cohort of n=15 mice. Please note that the motor network, the striatum network and the striato-motor network are highly correlated, thus we considered only the latter for the analysis and reduced the number of RSNs to 13. We performed a dual regression approach^110^ as described by^52^. With this approach, we derived an index of coupling strength (i.e. temporal synchronicity) of the voxels within each RSN, by averaging the Z-scores from the group-mean RSNs masks (thresholded at the 75^th^ percentile). Group level statistics were performed in SPSS v22 using a Linear Mixed-Model, with the fixed factors DREADD-Group and Time (2-levels, repeated measure), and with the individual mice as the random factor. Corresponding contrasts were used for post-hoc pairwise comparisons (LSD). P-values were considered significant at *p*<0.05 after False Discovery Rate (FDR) correction for multiple comparisons between RSNs.

### Ultra-high performance liquid chromatography (uHPLC)

To quantify norepinephrinergic (NE; epinephrine; MHPG), dopaminergic (DA; DOPAC; HVA), and serotonergic (5-HT; 5-HIAA) compounds, a reversed-phase uHPLC system coupled with electrochemical detection (RP-uHPLC-ECD) was used (Alexys^TM^ Neurotransmitter Analyzer, Antec Leyden, Zoeterwoude, Netherlands). In short, our previously validated RP-HPLC method with ion pairing chromatography was applied as described^111^, albeit with minor modifications regarding the installed column (BEH C18 Waters column, 150 mm x 1mm, 1.7μm particle size) and pump preference (LC110S pump, 497 bar; flow rate of 68μL/min), achieving the most optimal separation conditions in a RP-uHPLC setting. Levels of the monoamines and metabolites were calculated using Clarity software™ (DataApex Ltd., v6.2.0.208, 2015, Prague, Czech Republic).

Brain samples were defrosted to 4 °C and subsequently homogenized in cold 800μl sample buffer (50 mM citric acid, 50 mM phosphoric acid, 0.1 mM EDTA, 8 mM KCl and 1.8 mM octane-1-sulfonic acid sodium salt (OSA), adjusted to pH = 3.6), using a Bio-Gen PRO200 homogenizer (PRO Scientific Inc., Oxford, CT, USA; 60 s, 4°C). To remove excess proteins, 450 μl homogenate was transferred onto a 10,000 Da Amicon^®^ Ultra 0.5 Centrifugal Filter (Millipore, Ireland) that had been pre-washed twice using 450μl sample buffer (centrifugation: 14,000 × g, 20 min, 4 °C). The Amicon^®^ filter loaded with the homogenate was then centrifuged (14,000 × g, 20 min, 4°C). Finally, the filtrate was transferred into a polypropylene vial (0.3mL, Machery-Nagel GmbH & Co. KG, Germany) and automatically injected into the previously-mentioned uHPLC column by the Alexys™ AS110 sample injector.

### Statistics for behavior, pupillometry, immunohistochemistry and uHPLC

GraphPad Prism 8.0 was used for statistical analyses. We used independent samples t-tests when comparing two independent groups, and paired-samples t-tests when comparing the same group twice. When comparing more than two groups, we used one-way ANOVAs if there was a single independent variable, or two-way ANOVAs for two-factorial designs (e.g. region x treatment). Significant main effects and interactions were analyzed using Sidak’s post hoc tests.

### Gene expression

Gene expression data was obtained from the Allen Mouse Brain Atlas (AMBA)^50^ using the Allen Software Development Kit (SDK, stnava.github.io/ANTs/)^112^. Gene expression data in the AMBA is measured using in situ hybridization from: (i) sagittal section experiments with high genome coverage, and (ii) coronal section replications for approximately 3500 genes with restricted expression patterns in the brain (11). Transcriptional levels across a macroscopic cortical area were summarized as the ‘expression energy’ (the mean ISH intensity across voxels of that brain area)^50,113^.

## References

1. Sara, S. J. & Bouret, S. Orienting and Reorienting: The Locus Coeruleus Mediates Cognition through Arousal. Neuron 76, 130–141 (2012).

2. Manaye, K. F., McIntire, D. D., Mann, D. M. A. & German, D. C. Locus coeruleus cell loss in the aging human brain: A non-random process. J. Comp. Neurol. 358, 79–87 (1995).

3. McCall, J. G. et al. CRH Engagement of the Locus Coeruleus Noradrenergic System Mediates Stress-Induced Anxiety. Neuron 87, 605–620 (2015).

4. Naegeli, C. et al. Locus Coeruleus Activity Mediates Hyperresponsiveness in Posttraumatic Stress Disorder. Biol. Psychiatry 83, 254–262 (2018).

5. Isingrini, E. et al. Resilience to chronic stress is mediated by noradrenergic regulation of dopamine neurons. Nat. Neurosci. 19, 560–3 (2016).

6. Fortress, A. M. et al. Designer receptors enhance memory in a mouse model of Down syndrome. J. Neurosci. 35, 1343–53 (2015).

7. Bangasser, D. A., Eck, S. R. & Ordoñes Sanchez, E. Sex differences in stress reactivity in arousal and attention systems. Neuropsychopharmacology 44, 129–139 (2019).

8. Weinshenker, D. Long Road to Ruin: Noradrenergic Dysfunction in Neurodegenerative Disease. Trends Neurosci. 41, 211–223 (2018).

9. Carter, M. E. et al. Tuning arousal with optogenetic modulation of locus coeruleus neurons. Nat. Neurosci. 13, 1526–1533 (2010).

10. Takeuchi, T. et al. Locus coeruleus and dopaminergic consolidation of everyday memory. Nature 537, 357–362 (2016).

11. Kempadoo, K. A., Mosharov, E. V, Choi, S. J., Sulzer, D. & Kandel, E. R. Dopamine release from the locus coeruleus to the dorsal hippocampus promotes spatial learning and memory. Proc. Natl. Acad. Sci. U. S. A. 201616515 (2016). doi:10.1073/pnas.1616515114

12. Usher, M., Cohen, J. D., Servan-Schreiber, D., Rajkowski, J. & Aston-Jones, G. The role of locus coeruleus in the regulation of cognitive performance. Science (80-.). 283, 549–554 (1999).

13. Uematsu, A. et al. Modular organization of the brainstem noradrenaline system coordinates opposing learning states. Nat. Neurosci. 20, 1602–1611 (2017).

14. Li, L. et al. Stress Accelerates Defensive Responses to Looming in Mice and Involves a Locus Coeruleus-Superior Colliculus Projection. Curr. Biol. 28, 859–871.e5 (2018).

15. McCall, J. G. et al. Locus coeruleus to basolateral amygdala noradrenergic projections promote anxiety-like behavior. Elife 6, 1–23 (2017).

16. Hirschberg, S. et al. Functional dichotomy in spinal-vs prefrontal-projecting locus coeruleus modules splits descending noradrenergic analgesia from ascending aversion & anxiety in rats. Elife 1–26 (2017). doi:10.7554/eLife.29808

17. Borodovitsyna, O., Flamini, M. D. & Chandler, D. J. Acute Stress Persistently Alters Locus Coeruleus Function and Anxiety-like Behavior in Adolescent Rats. Neuroscience 373, 7–19 (2018).

18. Soya, S. et al. Orexin modulates behavioral fear expression through the locus coeruleus. Nat. Commun. 8, (2017).

19. Beas, B. S. et al. The locus coeruleus drives disinhibition in the midline thalamus via a dopaminergic mechanism. Nat. Neurosci. 21, 963–973 (2018).

20. Aston-Jones, G. & Cohen, J. D. An integrative theory of locus coeruleus-norepinephrine function: adaptive gain and optimal performance. Annu. Rev. Neurosci. 28, 403–50 (2005).

21. Bullmore, E. & Sporns, O. Complex brain networks: Graph theoretical analysis of structural and functional systems. Nat. Rev. Neurosci. 10, 186–198 (2009).

22. van den Heuvel, M. P. & Hulshoff Pol, H. E. Exploring the brain network: A review on resting-state fMRI functional connectivity. Eur. Neuropsychopharmacol. 20, 519–534 (2010).

23. Seeley, W. W. et al. Dissociable Intrinsic Connectivity Networks for Salience Processing and Executive Control. J. Neurosci. 27, 2349–2356 (2007).

24. C.W., B. & B.D., W. The locus coeruleus-noradrenergic system: Modulation of behavioral state and state-dependent cognitive processes. Brain Res. Rev. 42, 33–84 (2003).

25. Valentino, R. J. & Van Bockstaele, E. Convergent regulation of locus coeruleus activity as an adaptive response to stress. Eur. J. Pharmacol. 583, 194–203 (2008).

26. Arnsten, A. F. T. Stress signalling pathways that impair prefrontal cortex structure and function. Nat. Rev. Neurosci. 10, 410–422 (2009).

27. Corbetta, M., Patel, G. & Shulman, G. L. The Reorienting System of the Human Brain: From Environment to Theory of Mind. Neuron 58, 306–324 (2008).

28. Roeder, T. TYRAMINE AND OCTOPAMINE: Ruling Behavior and Metabolism. Annu. Rev. Entomol. 50, 447–477 (2005).

29. Hermans, E. J., Henckens, M. J. A. G., Joëls, M. & Fernández, G. Dynamic adaptation of large-scale brain networks in response to acute stressors. Trends Neurosci. 37, 304–14 (2014).

30. van Marle, H. J. F., Hermans, E. J., Qin, S. & Fernández, G. Enhanced resting-state connectivity of amygdala in the immediate aftermath of acute psychological stress. Neuroimage 53, 348–354 (2010).

31. Hermans, E. J. et al. Stress-related noradrenergic activity prompts large-scale neural network reconfiguration. Science 334, 1151–3 (2011).

32. Armbruster, B. N., Li, X., Pausch, M. H., Herlitze, S. & Roth, B. L. Evolving the lock to fit the key to create a family of G protein-coupled receptors potently activated by an inert ligand. Proc. Natl. Acad. Sci. 104, 5163–5168 (2007).

33. Parlato, R., Otto, C., Begus, Y., Stotz, S. & Schütz, G. Specific ablation of the transcription factor CREB in sympathetic neurons surprisingly protects against developmentally regulated apoptosis. Development 134, 1663–1670 (2007).

34. Roth, B. L. DREADDs for Neuroscientists. Neuron 89, 683–694 (2016).

35. Murphy, P. R., O’Connell, R. G., O’Sullivan, M., Robertson, I. H. & Balsters, J. H. Pupil diameter covaries with BOLD activity in human locus coeruleus. Hum. Brain Mapp. 35, 4140–4154 (2014).

36. Liu, Y., Rodenkirch, C., Moskowitz, N., Schriver, B. & Wang, Q. Dynamic Lateralization of Pupil Dilation Evoked by Locus Coeruleus Activation Results from Sympathetic, Not Parasympathetic, Contributions. Cell Rep. 20, 3099–3112 (2017).

37. Reimer, J. et al. Pupil fluctuations track rapid changes in adrenergic and cholinergic activity in cortex. Nat. Commun. 7, 13289 (2016).

38. Gomez, J. L. et al. Chemogenetics revealed: DREADD occupancy and activation via converted clozapine. Science (80-.). 503, 503–507 (2017).

39. Sturman, O., Germain, P.-L. & Bohacek, J. Exploratory rearing: a context- and stress-sensitive behavior recorded in the open-field test. Stress 21, 443–452 (2018).

40. Markicevic, M. et al. Cortical excitation:inhibition imbalance causes network specific functional hypoconnectivity: a DREADD-fMRI study. bioRxiv 492108 (2018). doi:10.1101/492108

41. Aksenov, D. P., Li, L., Miller, M. J., Iordanescu, G. & Wyrwicz, A. M. Effects of Anesthesia on BOLD Signal and Neuronal Activity in the Somatosensory Cortex. J. Cereb. Blood Flow Metab. 35, 1819–1826 (2015).

42. Jorm, C. M. & Stamford, J. A. Actions of the hypnotic anaesthetic, dexmedetomidine, on noradrenaline release and cell firing in rat locus coeruleus slices. Br. J. Anaesth. 71, 447–449 (1993).

43. Lakhlani, P. P. et al. Substitution of a mutant α2a-adrenergic receptor via ‘hit and run’ gene targeting reveals the role of this subtype in sedative, analgesic, and anesthetic-sparing responses in vivo. Proc Natl Acad Sci U S A. 94, 9950–9955 (1997).

44. Grandjean, J., Schroeter, A., Batata, I. & Rudin, M. NeuroImage Optimization of anesthesia protocol for resting-state fMRI in mice based on differential effects of anesthetics on functional connectivity patterns. Neuroimage 102, 838–847 (2014).

45. Bukhari, Q., Schroeter, A. & Rudin, M. Increasing isoflurane dose reduces homotopic correlation and functional segregation of brain networks in mice as revealed by resting-state fMRI. Sci. Rep. 8, 1–12 (2018).

46. Aston-Jones, G. Locus Coeruleus, A5 and A7 Noradrenergic Cell Groups. Rat Nerv. Syst. 259–294 (2004). doi:10.1016/B978-012547638-6/50012-2

47. Schwarz, L. A. & Luo, L. Organization of the locus coeruleus-norepinephrine system. Curr. Biol. 25, R1051–R1056 (2015).

48. Bouret, S. & Sara, S. J. Locus coeruleus activation modulates firing rate and temporal organization of odour-induced single-cell responses in rat piriform cortex. Eur. J. Neurosci. 16, 2371–2382 (2002).

49. Crick, F. C. & Koch, C. What is the function of the claustrum? Philos. Trans. R. Soc. B Biol. Sci. 360, 1271–1279 (2005).

50. Lein, E. S. et al. Genome-wide atlas of gene expression in the adult mouse brain. Nature 445, 168–76 (2007).

51. Filippini, N. et al. Distinct patterns of brain activity in young carriers of the APOE-4 allele. Proc. Natl. Acad. Sci. 106, 7209–7214 (2009).

52. Zerbi, V. et al. Dysfunctional Autism Risk Genes Cause Circuit-Specific Connectivity Deficits With Distinct Developmental Trajectories. Cereb. Cortex 28, 2495–2506 (2018).

53. Smith, C. C. & Greene, R. W. CNS Dopamine Transmission Mediated by Noradrenergic Innervation. J. Neurosci. 32, 6072–6080 (2012).

54. Models, A. Neuroanatomical Phenotyping in the Mouse: The Dopaminergic System. Vet. Pathol. 773, 753–773 (2005).

55. Schroeter, S. et al. Immunolocalization of the Cocaine- and l-Norepinephrine Transporter. 232, 211–232 (2000).

56. Mulvey, B. et al. Molecular and Functional Sex Differences of Noradrenergic Neurons in the Mouse Locus Coeruleus. Cell Rep. 23, 2225–2235 (2018).

57. Tervo, D. G. R. et al. A Designer AAV Variant Permits Efficient Retrograde Access to Projection Neurons. Neuron 92, 372–382 (2016).

58. Avery, M. C. & Krichmar, J. L. Neuromodulatory Systems and Their Interactions: A Review of Models, Theories, and Experiments. Front. Neural Circuits 11, 1–18 (2017).

59. Berton, O. & Nestler, E. J. New approaches to antidepressant drug discovery: beyond monoamines. Nat. Rev. Neurosci. 7, 137–51 (2006).

60. Lee, S. H. & Dan, Y. Neuromodulation of Brain States. Neuron 76, 109–222 (2012).

61. Lohani, S., Poplawsky, A. J., Kim, S.G. & Moghaddam, B. Unexpected global impact of VTA dopamine neuron activation as measured by opto-fMRI. Mol. Psychiatry 22, 585–594 (2017).

62. Roelofs, T. J. M. et al. A novel approach to map induced activation of neuronal networks using chemogenetics and functional neuroimaging in rats: A proof-of-concept study on the mesocorticolimbic system. Neuroimage 156, 109–118 (2017).

63. Giorgi, A. et al. Brain-wide Mapping of Endogenous Serotonergic Transmission via Chemogenetic fMRI. Cell Rep. 21, 910–918 (2017).

64. Grayson, D. S. et al. The Rhesus Monkey Connectome Predicts Disrupted Functional Networks Resulting from Pharmacogenetic Inactivation of the Amygdala. Neuron 91, 453–466 (2016).

65. Grandjean, J., Zerbi, V., Balsters, J., Wenderoth, N. & Rudina, M. The structural basis of large-scale functional connectivity in the mouse. J. Neurosci. 37, 0438–17 (2017).

66. Fox, M. D. & Greicius, M. Clinical applications of resting state functional connectivity. 4, (2010).

67. Totah, N. K., Neves, R. M., Panzeri, S., Logothetis, N. K. & Eschenko, O. The Locus Coeruleus Is a Complex and Differentiated Neuromodulatory System. Neuron 99, 1055–1068.e6 (2018).

68. Chandler, D. J., Gao, W. & Waterhouse, B. D. Heterogeneous organization of the locus coeruleus projections to prefrontal and motor cortices. 2014, 1–6 (2014).

69. Totah, N. K. B., Logothetis, N. K. & Eschenko, O. Noradrenergic ensemble-based modulation of cognition over multiple timescales. Brain Res. (2018). doi:10.1016/j.brainres.2018.12.031

70. Eschenko, O. et al. Tracing of noradrenergic projections using manganese-enhanced MRI. Neuroimage 59, 3252–3265 (2012).

71. Berridge, C. W., Isaac, S. O. & España, R. A. Additive wake-promoting actions of medial basal forebrain noradrenergic or α1- and β-receptor stimulation. Behav. Neurosci. 117, 350–359 (2003).

72. Manns, I. D., Lee, M. G., Modirrousta, M., Hou, Y. P. & Jones, B. E. Alpha 2 adrenergic receptors on GABAergic, putative sleep-promoting basal forebrain neurons. Eur. J. Neurosci. 18, 723–727 (2003).

73. Ramos, B. P. & Arnsten, A. F. T. Adrenergic pharmacology and cognition : Focus on the prefrontal cortex. 113, 523–536 (2007).

74. Ramos, B. P. et al. The beta-1 adrenergic antagonist, betaxolol, improves working memory performance in rats and monkeys. Biol. Psychiatry 58, 894–900 (2005).

75. Ramos, B. P., Colgan, L. A., Nou, E. & Arnsten, A. F. T. B2 Adrenergic Agonist, Clenbuterol, Enhances Working Memory Performance in Aging Animals. Neurobiol. Aging 29, 1060–1069 (2008).

76. Schwarz, L. A. et al. Viral-genetic tracing of the input-output organization of a central noradrenaline circuit. Nature 524, 88–92 (2015).

77. Kebschull, J. M. et al. High-Throughput Mapping of Single-Neuron Projections by Sequencing of Barcoded RNA NeuroResource High-Throughput Mapping of Single-Neuron Projections by Sequencing of Barcoded RNA. Neuron 91, 975–987 (2016).

78. Borodovitsyna, O., Joshi, N. & Chandler, D. Persistent Stress-Induced Neuroplastic Changes in the Locus Coeruleus/Norepinephrine System. Neural Plast. 2018, 1–14 (2018).

79. Vaisvaser, S. et al. Neural traces of stress: cortisol related sustained enhancement of amygdala-hippocampal functional connectivity. Front. Hum. Neurosci. 7, 1–11 (2013).

80. Seo, D. et al. Sex differences in neural responses to stress and alcohol context cues. Hum. Brain Mapp. 32, 1998–2013 (2011).

81. Sinha, R., Lacadie, C., Skudlarski, P. & Wexler, B. E. Neural circuits underlying emotional distress in humans. Ann. N. Y. Acad. Sci. 1032, 254–257 (2004).

82. van Oort, J. et al. How the brain connects in response to acute stress: A review at the human brain systems level. Neurosci. Biobehav. Rev. 83, 281–297 (2017).

83. Joëls, M. & Baram, T. Z. The neuro-symphony of stress. Nat. Rev. Neurosci. 10, 459–66 (2009).

84. Mana, M. J. & Grace, A. A. Chronic cold stress alters the basal and evoked electrophysiological activity of rat locus coeruleus neurons. Neuroscience 81, 1055–1064 (1997).

85. Pietrzak, R. H. et al. Association of Posttraumatic Stress Disorder With Reduced In Vivo Norepinephrine Transporter Availability in the Locus Coeruleus. JAMA Psychiatry 70, 1199 (2013).

86. Grandjean, J. et al. Chronic psychosocial stress in mice leads to changes in brain functional connectivity and metabolite levels comparable to human depression. Neuroimage 142, 544–552 (2016).

87. Magalhães, R. et al. The dynamics of stress : a longitudinal MRI study of rat brain structure and connectome. 1998–2006 (2018). doi:10.1038/mp.2017.244

88. Liberzon, I. & Phan, K. L. Brain-imaging studies of posttraumatic stress disorder. CNS Spectr. 8, 641–50 (2003).

89. Fitzgerald, J. M., DiGangi, J. A. & Phan, K. L. Functional Neuroanatomy of Emotion and Its Regulation in PTSD. Harv. Rev. Psychiatry 26, 116–128

90. Bangasser, D. A. & Valentino, R. J. Sex differences in stress-related psychiatric disorders: Neurobiological perspectives. Front. Neuroendocrinol. 35, 303–319 (2014).

91. Kawahara, H., Kawahara, Y. & Westerink, B. H. C. The noradrenaline-dopamine interaction in the rat medial prefrontal cortex studied by multi-probe microdialysis. Eur. J. Pharmacol. 418, 177–186 (2001).

92. Devoto, P., Flore, G., Saba, P., Fà, M. & Gessa, G. L. Stimulation of the locus coeruleus elicits noradrenaline and dopamine release in the medial prefrontal and parietal cortex. J. Neurochem. 92, 368–374 (2005).

93. Devoto, P., Flore, G., Saba, P., Fà, M. & Gessa, G. L. Co-release of noradrenaline and dopamine in the cerebral cortex elicited by single train and repeated train stimulation of the locus coeruleus. BMC Neurosci. 6, 1–11 (2005).

94. Antelman, S. & Caggiula, A. Norepinephrine-dopamine interactions and behavior. Science (80-.). 195, 646–653 (1977).

95. Swanson, L. W. & Hartman, B. K. The central adrenergic system. An immunofluorescence study of the location of cell bodies and their efferent connections in the rat utilizing dopamine-B-hydroxylase as a marker. J. Comp. Neurol. 163, 467–505 (1975).

96. Schallert, T., Whishaw, I. Q., Ramirez, V. D. & Teitelbaum, P. 6-hydroxydopamine and anticholinergic drugs. Science 202, 1216–7 (1978).

97. Ihalainen, J. A., Riekkinen, P. & Feenstra, M. G. P. Comparison of dopamine and noradrenaline release in mouse prefrontal cortex, striatum and hippocampus using microdialysis. Neurosci. Lett. 277, 71–74 (1999).

98. Hara, M. et al. Role of adrenoceptors in the regulation of dopamine/DARPP-32 signaling in neostriatal neurons. J. Neurochem. 113, 1046–1059 (2010).

99. Nicholas, A. P., Hökfelt, T. & Pieribone, V. A. The distribution and significance of CNS adrenoceptors examined with in situ hybridization. Trends Pharmacol. Sci. 17, 245–255 (1996).

100. Paschalis, A. et al. β1-Adrenoceptor distribution in the rat brain: An immunohistochemical study. Neurosci. Lett. 458, 84–88 (2009).

101. Pisani, A. et al. Activation of beta1-adrenoceptors excites striatal cholinergic interneurons through a cAMP-dependent, protein kinase-independent pathway. J. Neurosci. 23, 5272–5282 (2003).

102. Rommelfanger, K. S., Mitrano, D. A., Smith, Y. & Weinshenker, D. Light and electron microscopic localization of alpha-1 adrenergic receptor immunoreactivity in the rat striatum and ventral midbrain. Neuroscience 158, 1530–1540 (2009).

103. Bylund, D. B. & Snyder, S. H. Beta adrenergic receptor binding in membrane preparations from mammalian brain. Mol. Pharmacol. 12, 568–80 (1976).

104. Palacios, J. & Kuhar, M. Beta-adrenergic-receptor localization by light microscopic autoradiography. Science (80-.). 208, 1378–1380 (1980).

105. Reisine, T. D., Nagy, J. I., Beaumont, K., Fibiger, H. C. & Yamamura, H. I. The localization of receptor binding sites in the substantia nigra and striatum of the rat. Brain Res. 177, 241–252 (1979).

106. Mason, S. T. & Fibiger, H. C. Regional topography within noradrenergic locus coeruleus as revealed by retrograde transport of horseradish peroxidase. J. Comp. Neurol. 187, 703–724 (1979).

107. Vermeiren, Y. & De Deyn, P. P. Targeting the norepinephrinergic system in Parkinson’s disease and related disorders: The locus coeruleus story. Neurochem. Int. 102, 22–32 (2017).

108. Bakker, R., Tiesinga, P. & Kötter, R. The Scalable Brain Atlas: Instant Web-Based Access to Public Brain Atlases and Related Content. Neuroinformatics 13, 353–366 (2015).

109. Zerbi, V., Grandjean, J., Rudin, M. & Wenderoth, N. Mapping the mouse brain with rs-fMRI: An optimized pipeline for functional network identification. Neuroimage 123, 11–21 (2015).

110. Zuo, X. N. et al. Reliable intrinsic connectivity networks: Test-retest evaluation using ICA and dual regression approach. Neuroimage 49, 2163–2177 (2010).

111. Van Dam, D., Vermeiren, Y., Aerts, T. & De Deyn, P. P. Novel and sensitive reversed-phase high-pressure liquid chromatography method with electrochemical detection for the simultaneous and fast determination of eight biogenic amines and metabolites in human brain tissue. J. Chromatogr. A 1353, 28–39 (2014).

112. Fulcher, B. D., Murray, J. D., Zerbi, V. & Wang, X.-J. Multimodal gradients across mouse cortex. 1–7 (2018). doi:10.1101/393215

113. Ng, L. et al. An anatomic gene expression atlas of the adult mouse brain. Nat. Neurosci. 12, 356–362 (2009).

