## Supplementary Figures for "Rapid Reconfiguration of the Functional Connectome after Chemogenetic Locus Coeruleus Activation"

**Valerio Zerbi[1,7*#], Amalia Floriou-Servou[2,7*], Marija Markicevic[1,7], Yannick Vermeiren[3,5], Oliver Sturman[2,7], Mattia Privitera[2,7], Lukas von Ziegler[2,7], Kim David Ferrari[4,7], Bruno Weber[4,7], Peter Paul De Deyn[3,5,6], Nici Wenderoth[1,7#], Johannes Bohacek[2,7#]**

[1] Neural Control of Movement Lab, Department of Health Sciences and Technology, ETH Zürich, Switzerland

[2] Laboratory of Molecular and Behavioral Neuroscience, Institute for Neuroscience, Department of Health Sciences and Technology, ETH Zurich, Switzerland

[3] Department of Biomedical Sciences, Laboratory of Neurochemistry and Behavior, Institute Born-Bunge, University of Antwerp, Wilrijk (Antwerp), Belgium

[4] Experimental Imaging and Neuroenergetics, Institute of Pharmacology and Toxicology, University of Zurich, Switzerland

[5] Department of Neurology and Alzheimer Research Center, University of Groningen and University Medical Center Groningen (UMCG), Groningen, Netherlands

[6] Department of Neurology, Memory Clinic of Hospital Network Antwerp (ZNA) Middelheim and Hoge Beuken, Antwerp, Belgium

[7] Neuroscience Center Zurich, ETH Zurich and University of Zurich, Switzerland

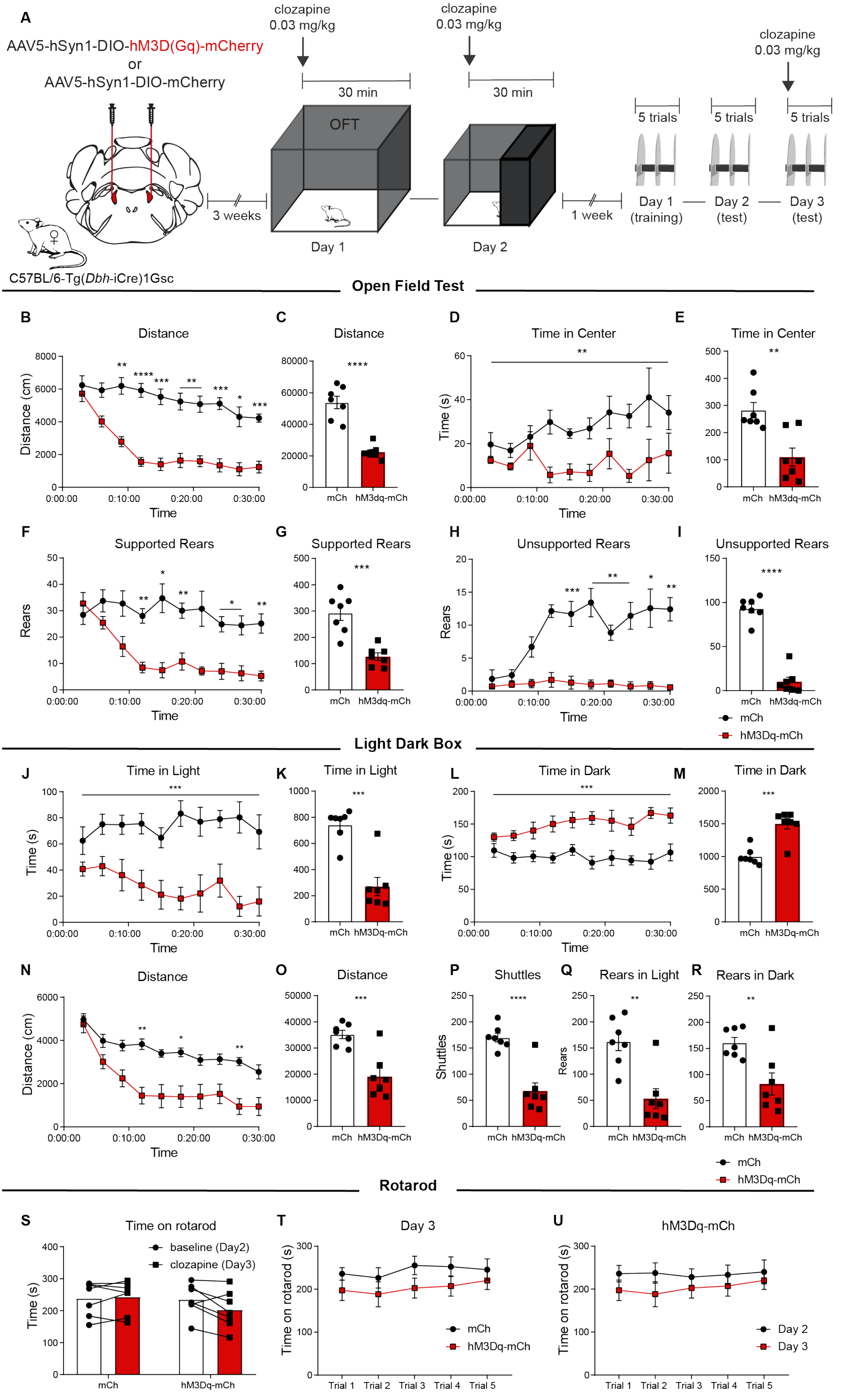
A
AAV5-hSyn1-DIO-hM3D(Gq)-mCherry
or
AAV5-hSyn1-DIO-mCherry
C57BL/6-Tg(Dbh-iCre)1Gsc
3 weeks
clozapine
0.03 mg/kg
30 min
OFT
Day 1
Day 2
1 week
5 trials
Day 1
(training)
Day 2
(test)
Day 3
(test)
Open Field Test
B Distance
C Distance
D Time in Center
E Time in Center
F Supported Rears
G Supported Rears
H Unsupported Rears
I Unsupported Rears
Distance (cm)
Time (s)
Rears
Time
mCh
hM3dq-mCh
hM3Dq-mCh
Light Dark Box
J Time in Light
K Time in Light
L Time in Dark
M Time in Dark
N Distance
O Distance
P Shuttles
Q Rears in Light
R Rears in Dark
Shuttles
Rotarod
S Time on rotarod
baseline (Day2)
clozapine (Day3)
T Day 3
U hM3Dq-mCh
Time on rotarod (s)
Trial 1
Trial 2
Trial 3
Trial 4
Trial 5
Day 2
Day 3

**Figure S1.** (**A**) Diagram showing the time-course of behavioral tests after virus delivery in the LC of female DBH-iCre mice. (**B-I**) Mice were placed in the OFT directly after clozapine injection for 30 minutes. Mice expressing hM3Dq-mCh travelled less distance compared to mice expressing only mCh (**B, C:** main effect of group: $F(1,12)=53.71$, $p<0.0001$, interaction: $F(9,108)=6.32$, $p<0.0001$, two-way ANOVA with Sidak *post hoc* tests), spent less time in the center (**D, E:** main effect of group $F(1,12)=15.81$, $p=0.0018$, two-way ANOVA) and performed fewer supported rears (**F, G:** main effect of group $F(1,12)=27.46$, $p=0.0002$, interaction: $F(9,108)=4.24$, $p=0.0001$, two-way ANOVA with Sidak *post hoc* tests) and unsupported rears (**H, I:** main effect of group: $F(1,12)=137.9$, $p<0.0001$, interaction: $F(9,108)=5.26$, $p<0.0001$, two-way ANOVA with Sidak *post hoc* tests). (**J-R**) Mice were placed in the light dark box directly after clozapine injection for 30 minutes. Mice expressing hM3Dq-mCh spent less time in the light compartment in comparison to mCh controls (**J, K:** main effect of group: $F(1,12)=31.44$, $p=0.0001$, two-way ANOVA), more time in the dark compartment (**L, M:** main effect of group: $F(1,12)=28.75$, $p=0.0002$, two-way ANOVA), and travelled less distance (**N, O:** main effect of group: $F(1,12)=20.65$, $p=0.0007$, interaction: $F(9,108)=2.67$, $p=0.0072$, two-way ANOVA with Sidak *post hoc* tests). (**P-R**) Compared to mCh controls, hM3Dq-mCh mice also performed fewer shuttles between the light and the dark compartment (**P:** $t(12)=5.81$, $p<0.0001$, unpaired t test) and fewer rears in both compartments (**Q:** $t(12)=4.26$, $p=0.0011$, **R:** $t(12)=3.36$, $p=0.0057$, unpaired t test). (**S-U**) To assess gross motor function we trained the same mice on the Rotarod (Day 1, 5 trials) and tested them at baseline (Day 2, 5 trials) and immediately after clozapine injection (Day 3, 5 trials). The average performance from all trials of hM3Dq-mCh mice was slightly lower, not significantly different from mCh controls (**S:** main effect of group: $F(1,12)=2.61$, $p=0.1318$, interaction virus x group: $F(1,12)=4.61$, $p=0.0528$, two-way ANOVA). On Day 3, the performance of hM3Dq-mCh and mCh mice was not significantly different over the course of 5 trials (**T:** main effect of group: $F(1,12)=1.80$, $p=0.2040$, interaction trial x group: $F(4,48)=0.45$, $p=0.7743$, two-way ANOVA) and the performance of hM3Dq-mCh was not different between Day 2 and Day 3 over the course of 5 trials (**U:** main effect of group: $F(1,12)=1.18$, $p=0.2982$, interaction trial x group: $F(4,48)=0.46$, $p=0.7667$). *$p<0.05$, **$p<0.01$, ***$p<0.001$, ****$p<0.0001$. Data represent mean ± SEM

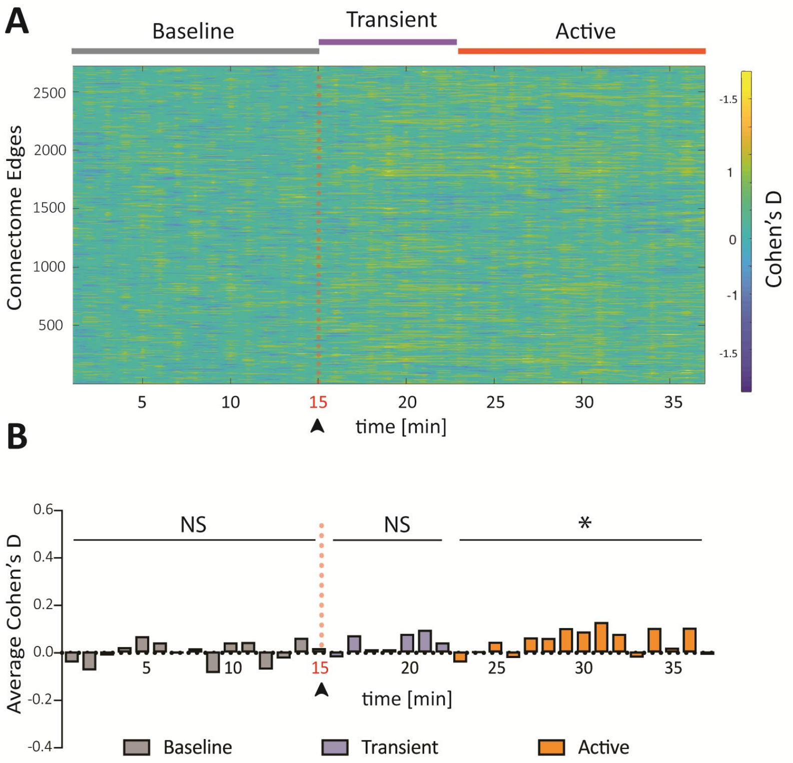


**Figure S2. Minor FC changes between anesthesia with 1% isoflurane versus anesthesia with 0.5% isoflurane + medetomidine.** Effect-size (Cohen's D) analysis of Functional Connectivity (FC) is shown for single-edges (n=2724) between mCh mice (n=7) under two anesthesia conditions (see methods) (**A**) and for the average across all edges (**B**). The distribution of the data reveals a reduction of connectivity in multiple edges in the "1% isoflurane"-condition compared to the "0.5% isoflurane + medetomidine"-condition, about 30 minutes after the start of the session (Wilcoxon two-tailed test: $p$=0.9780 for baseline period, $p$=0.0781 for transient period and $p$=0.0181 for active period). The average net-effect of anesthesia is approximately 7 times smaller than the effect of LC-NE DREADD activation.

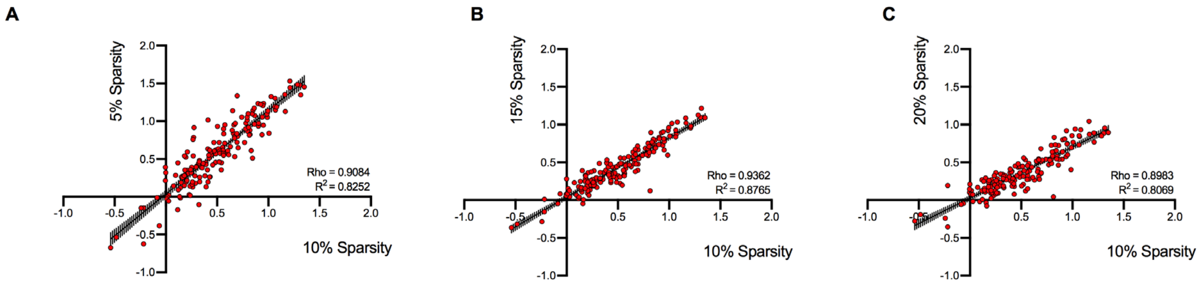


**Figure S3. The Node Modulation Index (NMI) effect size is a robust index and correlates at different connectome sparsity-thresholds.** The NMI effect size between mCh (n=7) and mCh-hM3Dq (n=11) under 1% isoflurane anesthesia is calculated at different sparsity thresholds (5%, 15%, 20%) of the connectome matrix. In each case, the resulting indices were highly linearly correlated (Pearson's correlation, $p$-value < 0.0001).

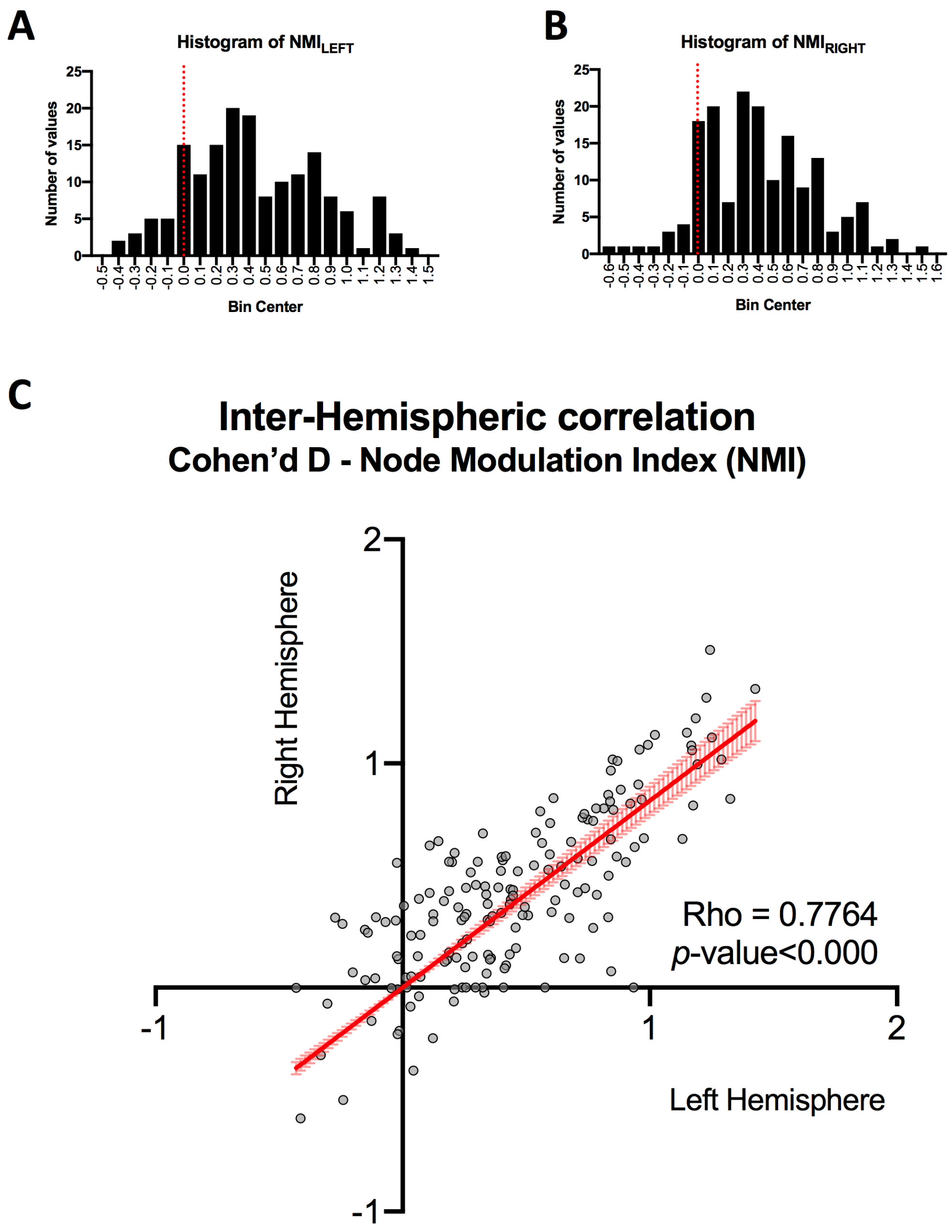


**Figure S4. Coherent FC changes in the left- and right- hemisphere after LC-NE activation.** (**A**) Right-skewed distribution plots of Node Modulation Index effect size (Cohen's D) in the left and in the right (**B**) hemispheres, demonstrating hyper-connectivity in both hemispheres. (**C**) The NMI measured in the left hemisphere shows positive linear correlation with respect to right hemisphere NMI (Pearson Rho=0.7764, p<0.0001). Data from mCh (n=7) and mCh-hM3Dq (n=11) under 1% isoflurane anesthesia.

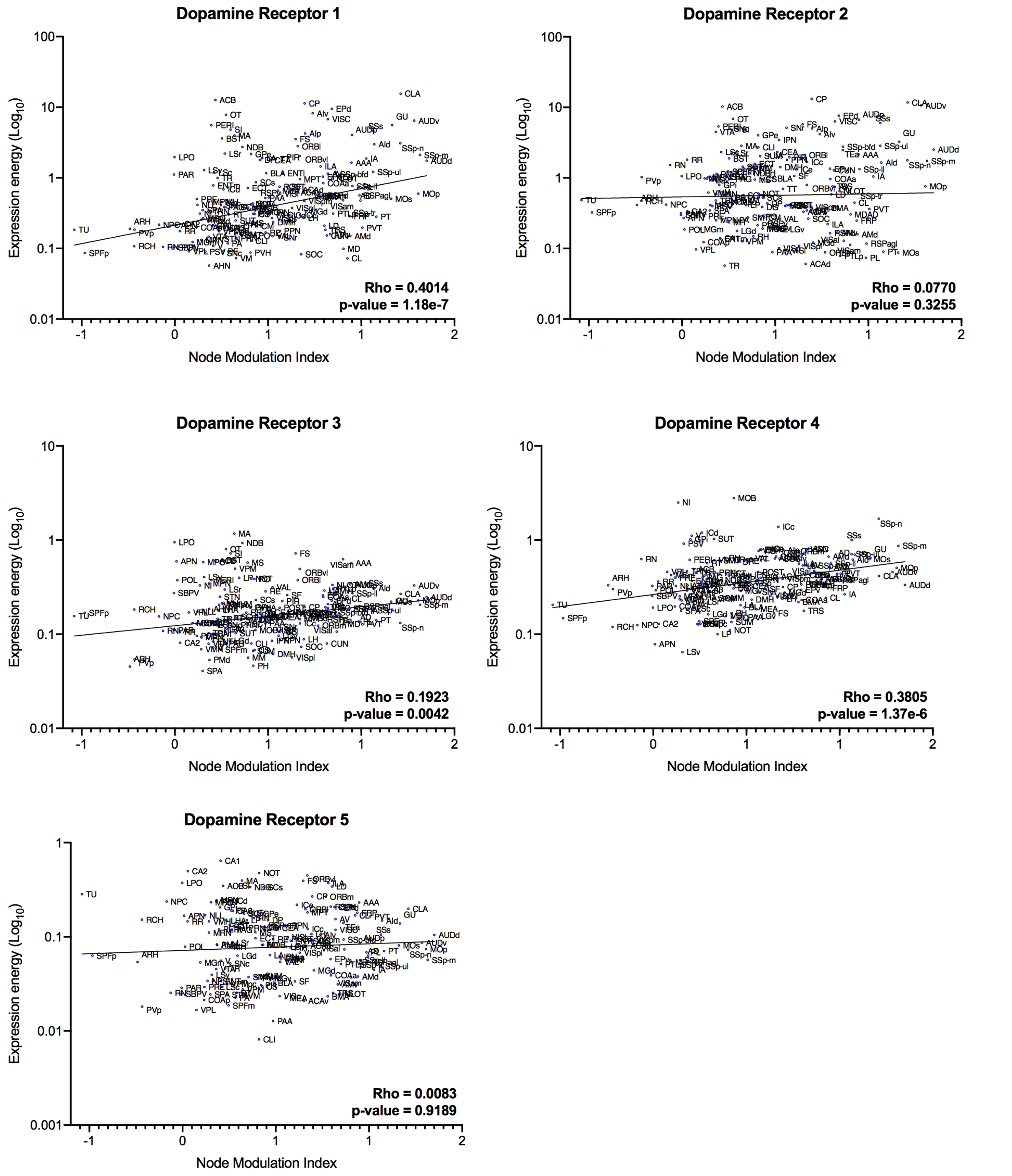


**Figure S5. Node Modulation Index maps correlate with D1 and D4 receptor gene-transcript maps.** Spearman correlation coefficients, Rho, and associated p-value (FDR corrected) between Node Modulation Index and the transcriptional maps of genes coding dopamine receptor subunits (D1,D2,D3,D4,D5).

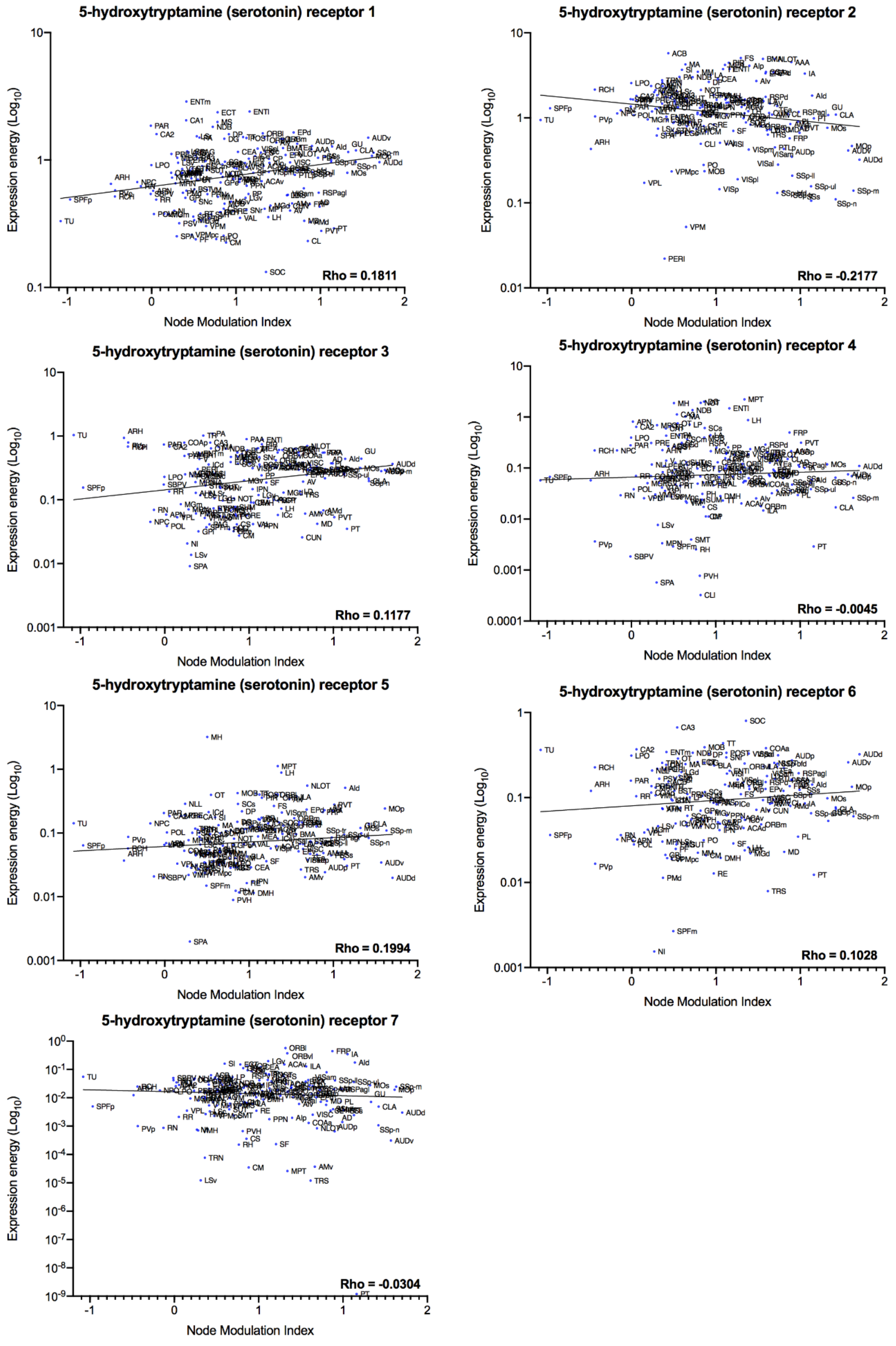


**Figure S6. Node Modulation Index maps do not correlate with serotonin receptor gene-transcript maps.** Spearman correlation coefficients, Rho, and associated p-value (FDR corrected) between Node Modulation Index and the transcriptional maps of genes coding serotonin receptor subunits (1,2,3,4,5,6,7). None of the transcriptional maps show significant correlation with the NMI.

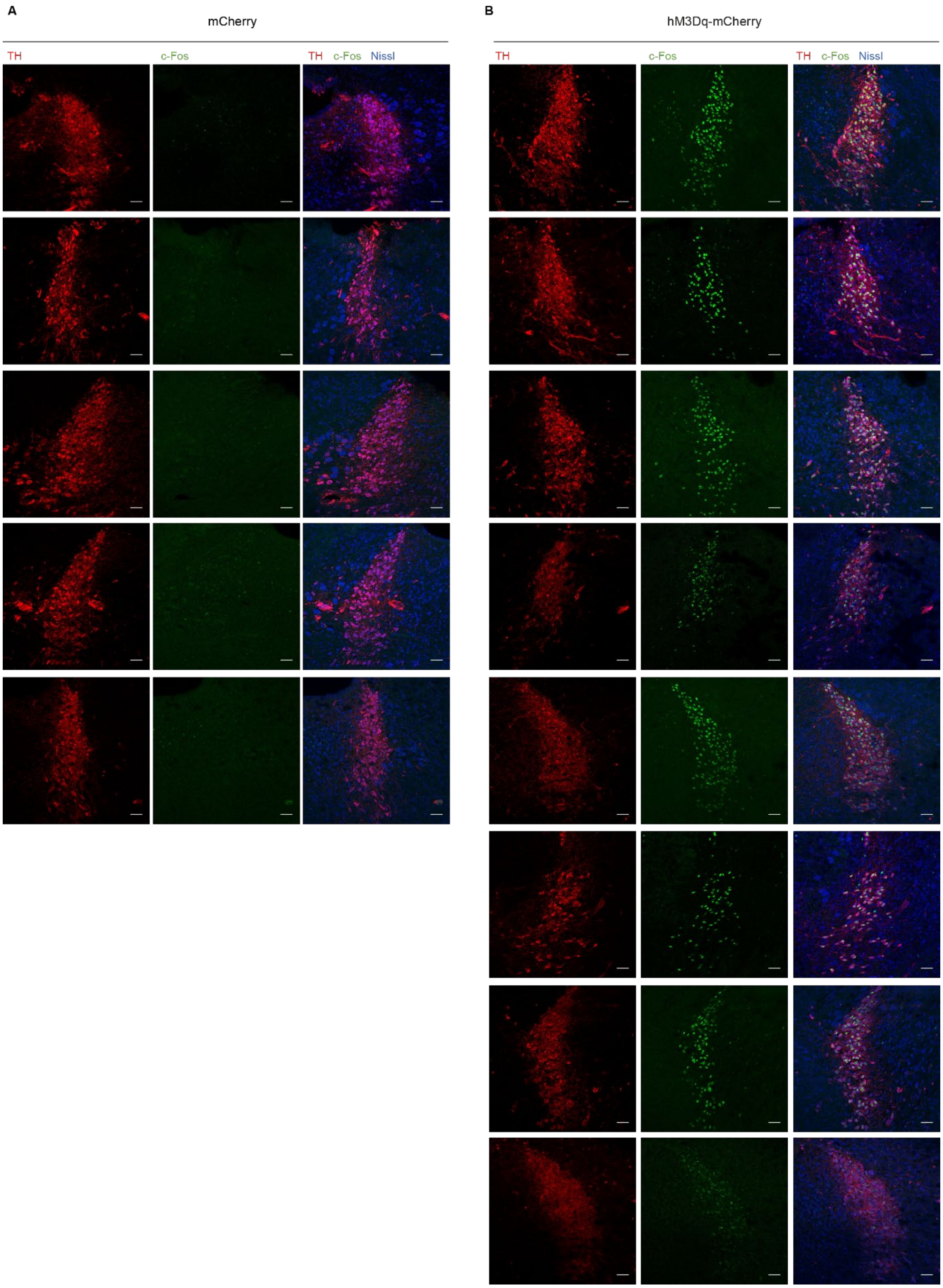


**Figure S7.** Representative LC images of every mouse used for the fMRI scans. Every row shows 3 images from one mouse, stained for TH (left), cFos (middle), and the merged picture including a Nissl stain (right). Tissue was collected 90 minutes after 0.03mg/kg clozapine injection, and cFos is activated in all mice expressing hM3Dq-mCh (B), but not in mCh controls (A). Tissue collected as described in Figure 5. All scale bars: 50 μm.

A

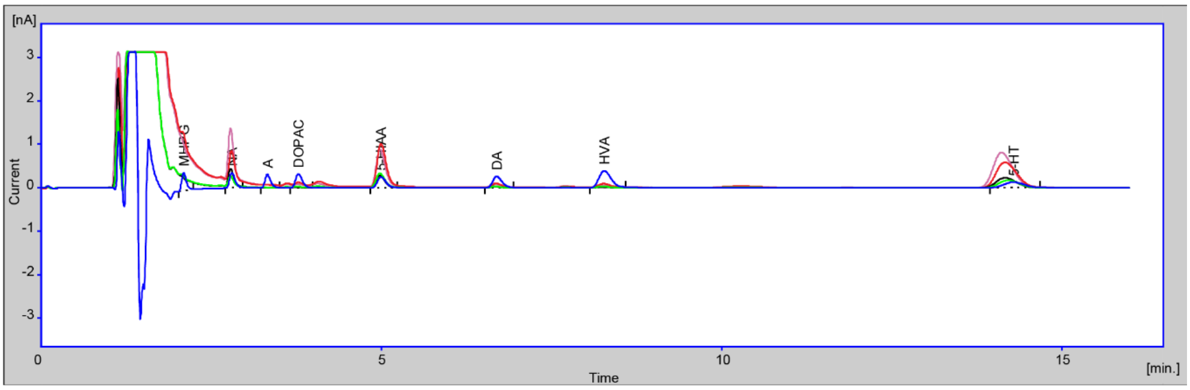


B

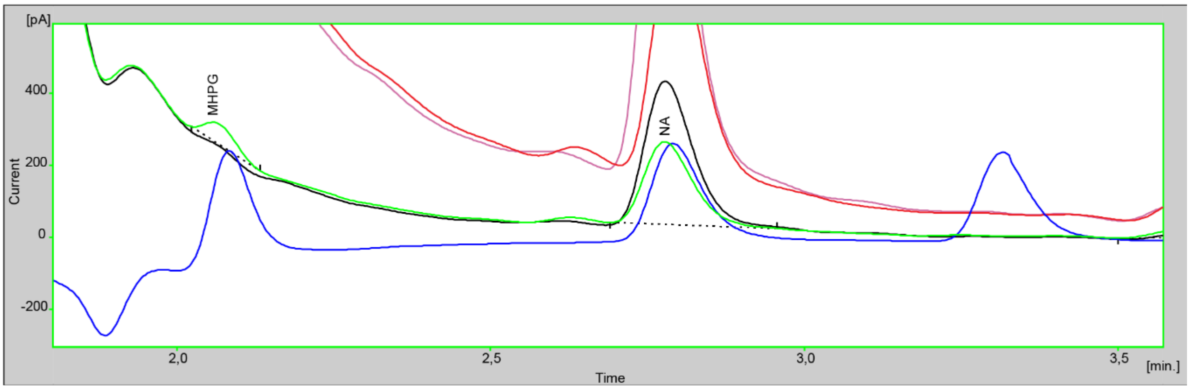


**Figure S8.** (**A**) uHPLC chromatograms of hippocampus samples from an mCh mouse (undiluted in pink, 3 times diluted in black), an hM3Dq-mCh mouse (undiluted in red, 3 times diluted in green) and the standard (blue). (**B**) Magnification of (A) showing the chromatogram from an hM3Dq-mCh mouse with higher MHPG and lower NA values (green) resulting in a high norepinephrine turnover ratio (MHPG/NE), in contrast with the chromatogram from an mCh mouse (black). The standard is in blue. Abbreviations: 5-HIAA: 5-hydroxyindoleacetic acid; 5-HT: 5-hydroxytryptamine (serotonin); A: adrenaline (epinephrine); DA: dopamine; DOPAC: 3,4-dihydroxyphenylacetic acid; HVA: homovanillic acid; MHPG: 3-methoxy-4-hydroxyphenylglycol; nA: nanoampere; NA: noradrenaline (norepinephrine); pA: picoampere.

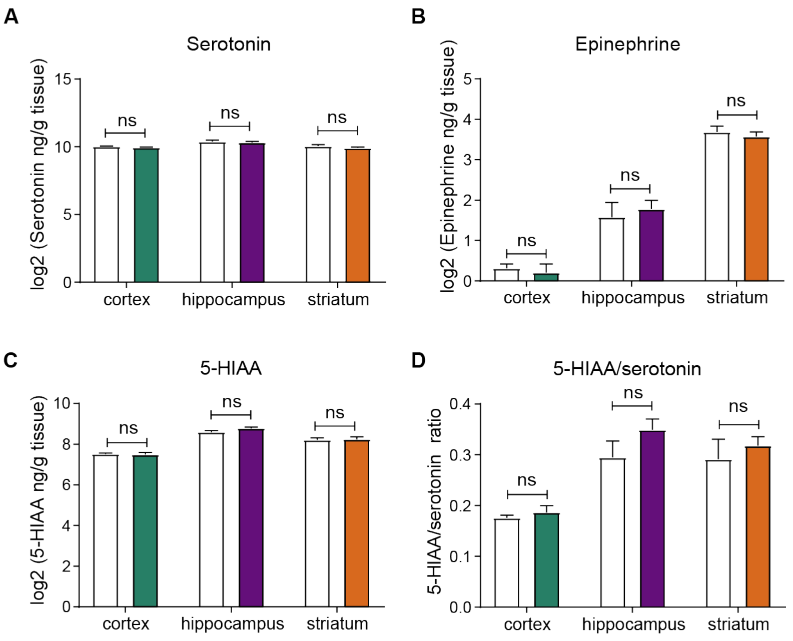


**Figure S9.** Levels of serotonin, epinephrine, 5-HIAA, and serotonin turnover (5-HIAA/serotonin ratio) unchanged in response to LC activation. Samples were collected 90 minutes after 0.03 mg/kg clozapine injection (two-way ANOVA), as explained in Figure 6A. Data represent mean ± SEM

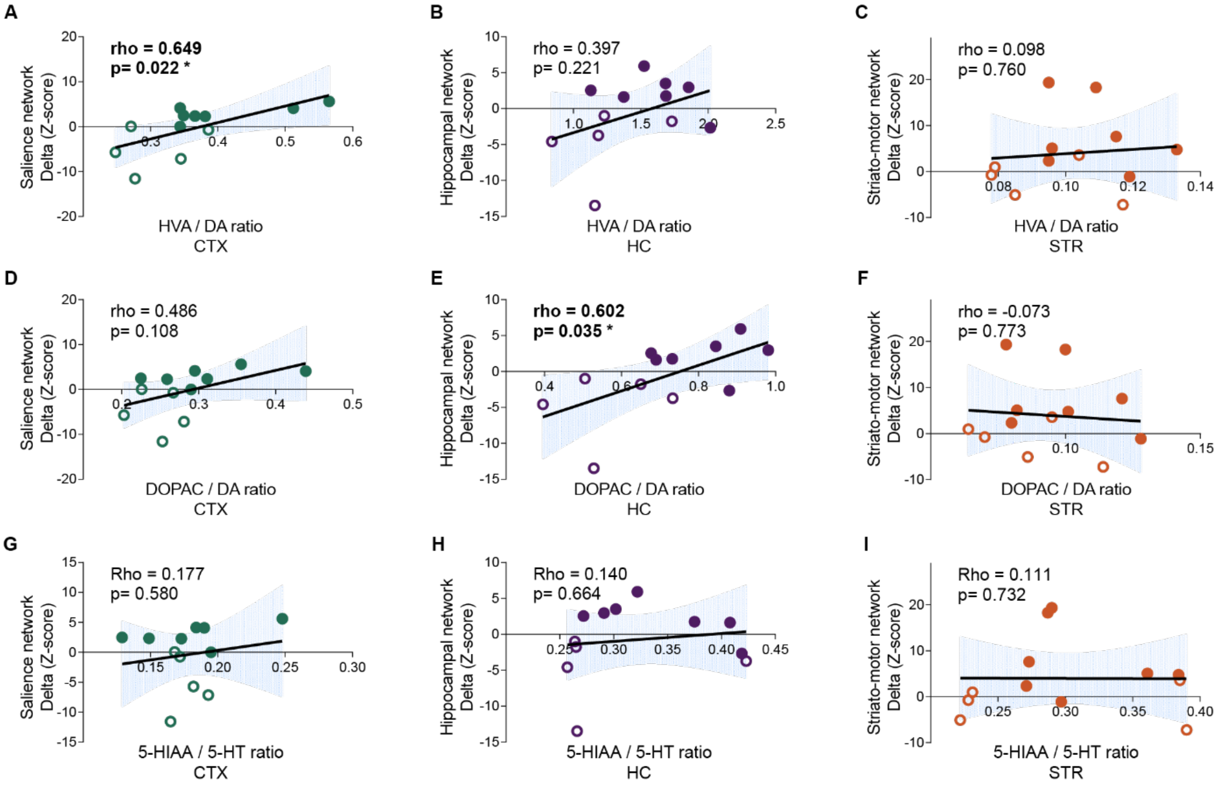


**Figure S10.** Spearman correlation coefficients Rho, and associated p-value (FDR corrected) between the turnover of DA (**A-F**) and 5-HT (**G-I**) in the cortex, hippocampus and striatum and the changes in network connectivity in the Salience Network (left), Hippocampal Network (middle), and Striato-Motor Networks (right).

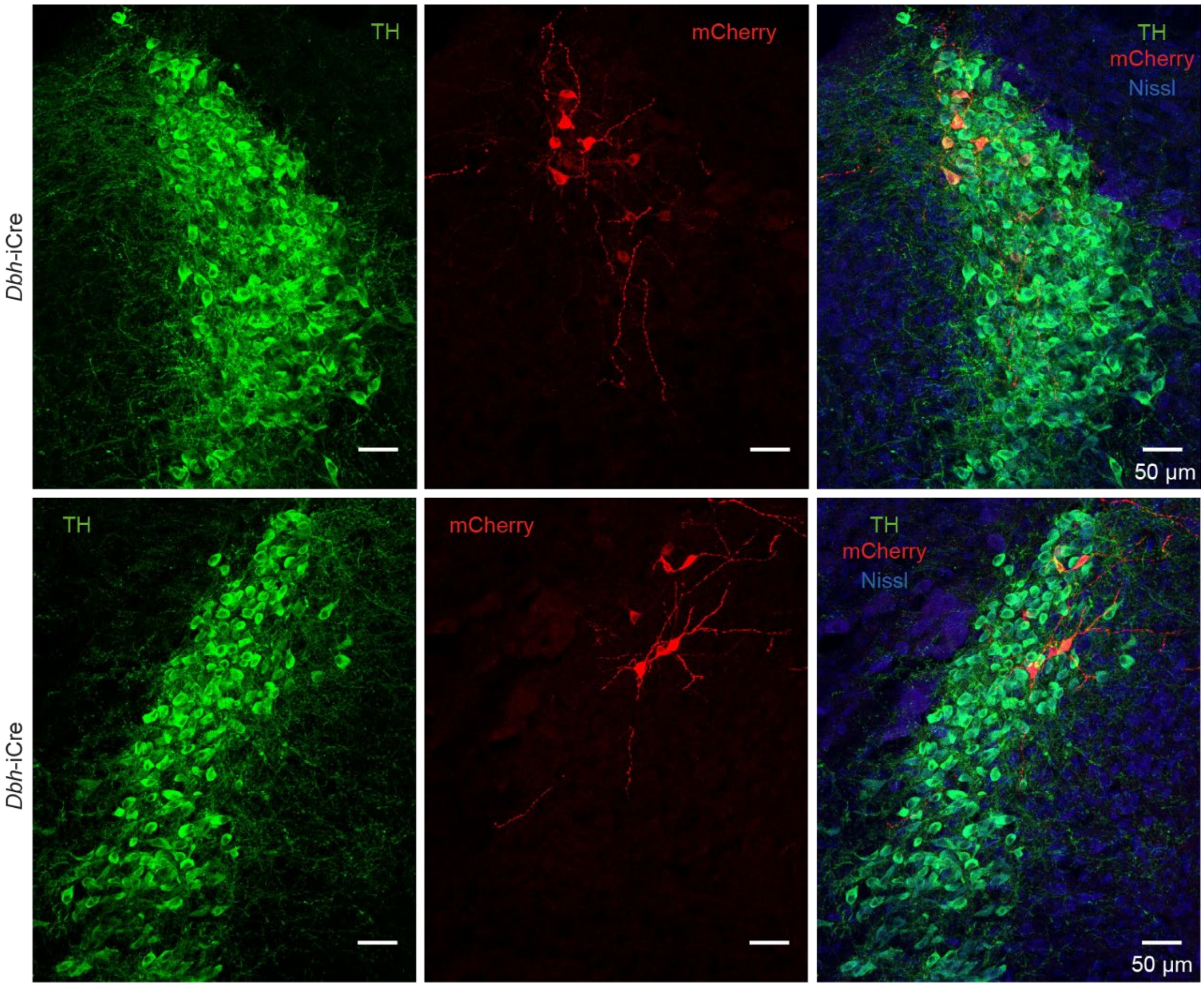


**Figure S11.** Representative images from 2 DBH-iCre animals showing mCherry+ neurons in the LC after delivery of a Cre-dependent, mCherry-expressing retro-AAV2 in the dorsolateral caudate putamen.
