## Supplementary Table 1 for "Rapid Reconfiguration of the Functional Connectome after Chemogenetic Locus Coeruleus Activation"

| Ranking | Node Modulation Index (Cohen's D) | Short Name | Full Name | Structure |
|---|---|---|---|---|
| 1 | 1,3511 | AUDd | 'Dorsal auditory area' | 'Isocortex' |
| 2 | 1,3154 | SSp-m | 'Primary somatosensory area, mouth' | 'Isocortex' |
| 3 | 1,3061 | MOp | 'Primary motor area' | 'Isocortex' |
| 4 | 1,2845 | AUDv | 'Ventral auditory area' | 'Isocortex' |
| 5 | 1,2113 | CLA | 'Claustrum' | 'Cortical Subplate' |
| 6 | 1,2094 | SSp-n | 'Primary somatosensory area, nose' | 'Isocortex' |
| 7 | 1,1659 | GU | 'Gustatory areas' | 'Isocortex' |
| 8 | 1,1608 | MOs | 'Secondary motor area' | 'Isocortex' |
| 9 | 1,0808 | PT | 'Parataenial nucleus' | 'Thalamus' |
| 10 | 1,0702 | AId | 'Agranular insular area, dorsal part' | 'Isocortex' |
| 11 | 1,0651 | SSs | 'Supplemental somatosensory area' | 'Isocortex' |
| 12 | 1,0640 | SSp-ul | 'Primary somatosensory area, upper limb' | 'Isocortex' |
| 13 | 1,0271 | IA | 'Intercalated amygdalar nucleus' | 'Striatum' |
| 14 | 1,0056 | PVT | 'Paraventricular nucleus of the thalamus' | 'Thalamus' |
| 15 | 0,9958 | AD | 'Anterodorsal nucleus' | 'Thalamus' |
| 16 | 0,9902 | RSPagl | 'Retrosplenial area, lateral agranular part' | 'Isocortex' |
| 17 | 0,9869 | PL | 'Prelimbic area' | 'Isocortex' |
| 18 | 0,9528 | SSp-ll | 'Primary somatosensory area, lower limb' | 'Isocortex' |
| 19 | 0,9501 | AUDp | 'Primary auditory area' | 'Isocortex' |
| 20 | 0,9471 | AAA | 'Anterior amygdalar area' | 'Striatum' |
| 21 | 0,9412 | AMd | 'Anteromedial nucleus, dorsal part' | 'Thalamus' |
| 22 | 0,9372 | FRP | 'Frontal pole, cerebral cortex' | 'Isocortex' |
| 23 | 0,9257 | CL | 'Central lateral nucleus of the thalamus' | 'Thalamus' |
| 24 | 0,9046 | MD | 'Mediodorsal nucleus of thalamus' | 'Thalamus' |
| 25 | 0,9027 | VISam | 'Anteromedial visual area' | 'Isocortex' |
| 26 | 0,9001 | SSp-tr | 'Primary somatosensory area, trunk' | 'Isocortex' |
| 27 | 0,8686 | VISal | 'Anterolateral visual area' | 'Isocortex' |
| 28 | 0,8641 | SSp-bfd | 'Primary somatosensory area, barrel field' | 'Isocortex' |
| 29 | 0,8628 | TEa | 'Temporal association areas' | 'Isocortex' |
| 30 | 0,8494 | PTLp | 'Posterior parietal association areas' | 'Isocortex' |
| 31 | 0,8455 | NLOT | 'Nucleus of the lateral olfactory tract' | 'Olfactory Areas' |
| 32 | 0,8419 | EPd | 'Endopiriform nucleus, dorsal part' | 'Cortical Subplate' |
| 33 | 0,8329 | AMv | 'Anteromedial nucleus, ventral part' | 'Thalamus' |
| 34 | 0,8207 | AV | 'Anteroventral nucleus of thalamus' | 'Thalamus' |
| 35 | 0,8205 | VISC | 'Visceral area' | 'Isocortex' |
| 36 | 0,8151 | CUN | 'Cuneiform nucleus' | 'Midbrain' |
| 37 | 0,8075 | TRS | 'Triangular nucleus of septum' | 'Pallidum' |
| 38 | 0,8023 | LD | 'Lateral dorsal nucleus of thalamus' | 'Thalamus' |
| 39 | 0,7960 | COAa | 'Cortical amygdalar area, anterior part' | 'Olfactory Areas' |
| 40 | 0,7947 | RSPd | 'Retrosplenial area, dorsal part' | 'Isocortex' |
| 41 | 0,7944 | EPv | 'Endopiriform nucleus, ventral part' | 'Cortical Subplate' |
| 42 | 0,7817 | ILA | 'Infralimbic area' | 'Isocortex' |
| 43 | 0,7786 | BMA | 'Basomedial amygdalar nucleus' | 'Cortical Subplate' |
| 44 | 0,7683 | ORBm | 'Orbital area, medial part' | 'Isocortex' |
| 45 | 0,7395 | AIv | 'Agranular insular area, ventral part' | 'Isocortex' |
| 46 | 0,7048 | MGd | 'Medial geniculate complex, dorsal part' | 'Thalamus' |
| 47 | 0,6984 | AIp | 'Agranular insular area, posterior part' | 'Isocortex' |
| 48 | 0,6964 | CP | 'Caudoputamen' | 'Striatum' |
| 49 | 0,6946 | VISpm | 'posteromedial visual area' | 'Isocortex' |
| 50 | 0,6913 | LH | 'Lateral habenula' | 'Thalamus' |
| 51 | 0,6776 | SOC | 'Superior olivary complex' | 'Pons' |
| 52 | 0,6714 | ICc | 'Inferior colliculus, central nucleus' | 'Midbrain' |
| 53 | 0,6708 | MPT | 'Medial pretectal area' | 'Midbrain' |
| 54 | 0,6695 | ORBvl | 'Orbital area, ventrolateral part' | 'Isocortex' |
| 55 | 0,6637 | ACAd | 'Anterior cingulate area, dorsal part' | 'Isocortex' |
| 56 | 0,6589 | ORBl | 'Orbital area, lateral part' | 'Isocortex' |
| 57 | 0,6523 | ACAv | 'Anterior cingulate area, ventral part' | 'Isocortex' |
| 58 | 0,6477 | FS | 'Fundus of striatum' | 'Striatum' |
| 59 | 0,6289 | VISpl | 'Posterolateral visual area' | 'Isocortex' |
| 60 | 0,6135 | ICe | 'Inferior colliculus, external nucleus' | 'Midbrain' |
| 61 | 0,6026 | SF | 'Septofimbrial nucleus' | 'Striatum' |
| 62 | 0,5809 | ENTl | 'Entorhinal area, lateral part' | 'Hippocampal Formation' |
| 63 | 0,5762 | PIR | 'Piriform area' | 'Olfactory Areas' |
| 64 | 0,5743 | PP | 'Peripeduncular nucleus' | 'Thalamus' |
| 65 | 0,5653 | VISl | 'Lateral visual area' | 'Isocortex' |
| 66 | 0,5637 | PPN | 'Pedunculopontine nucleus' | 'Midbrain' |
| 67 | 0,5622 | POST | 'Postsubiculum' | 'Hippocampal Formation' |
| 68 | 0,5615 | SNr | 'Substantia nigra, reticular part' | 'Midbrain' |
| 69 | 0,5566 | LGv | 'Ventral part of the lateral geniculate complex' | 'Thalamus' |
| 70 | 0,5532 | MEA | 'Medial amygdalar nucleus' | 'Striatum' |

| Ranking | Node Modulation Index (Cohen's D) | Short Name | Full Name | Structure |
|---|---|---|---|---|
| 71 | 0,5420 | TT | 'Taenia tecta' | 'Olfactory Areas' |
| 72 | 0,5415 | IMD | 'Intermediodorsal nucleus of the thalamus' | 'Thalamus' |
| 73 | 0,5278 | DMH | 'Dorsomedial nucleus of the hypothalamus' | 'Hypothalamus' |
| 74 | 0,5239 | VAL | 'Ventral anterior-lateral complex of the thalamus' | 'Thalamus' |
| 75 | 0,5208 | VISp | 'Primary visual area' | 'Isocortex' |
| 76 | 0,5206 | IPN | 'Interpeduncular nucleus' | 'Midbrain' |
| 77 | 0,5122 | CEA | 'Central amygdalar nucleus' | 'Striatum' |
| 78 | 0,5042 | RSPv | 'Retrosplenial area, ventral part' | 'Isocortex' |
| 79 | 0,4896 | BLA | 'Basolateral amygdalar nucleus' | 'Cortical Subplate' |
| 80 | 0,4871 | RE | 'Nucleus of reuniens' | 'Thalamus' |
| 81 | 0,4849 | PAA | 'Piriform-amygdalar area' | 'Olfactory Areas' |
| 82 | 0,4707 | MGv | 'Medial geniculate complex, ventral part' | 'Thalamus' |
| 83 | 0,4682 | LA | 'Lateral amygdalar nucleus' | 'Cortical Subplate' |
| 84 | 0,4576 | DP | 'Dorsal peduncular area' | 'Olfactory Areas' |
| 85 | 0,4416 | CM | 'Central medial nucleus of the thalamus' | 'Thalamus' |
| 86 | 0,4348 | SCs | 'Superior colliculus, sensory related' | 'Midbrain' |
| 87 | 0,4324 | MOB | 'Main olfactory bulb' | 'Olfactory Areas' |
| 88 | 0,4308 | DG | 'Dentate gyrus' | 'Hippocampal Formation' |
| 89 | 0,4278 | CS | 'Superior central nucleus raphe' | 'Pons' |
| 90 | 0,4266 | PO | 'Posterior complex of the thalamus' | 'Thalamus' |
| 91 | 0,4210 | PH | 'Posterior hypothalamic nucleus' | 'Hypothalamus' |
| 92 | 0,4193 | SUM | 'Supramammillary nucleus' | 'Midbrain' |
| 93 | 0,4108 | NOT | 'Nucleus of the optic tract' | 'Midbrain' |
| 94 | 0,4104 | CLI | 'Central linear nucleus raphe' | 'Midbrain' |
| 95 | 0,4088 | GPe | 'Globus pallidus, external segment' | 'Pallidum' |
| 96 | 0,4057 | PVH | 'Paraventricular hypothalamic nucleus' | 'Hypothalamus' |
| 97 | 0,3926 | MM | 'Medial mammillary nucleus' | 'Hypothalamus' |
| 98 | 0,3915 | ECT | 'Ectorhinal area' | 'Isocortex' |
| 99 | 0,3905 | MS | 'Medial septal nucleus' | 'Pallidum' |
| 100 | 0,3825 | RH | 'Rhomboid nucleus' | 'Thalamus' |
| 101 | 0,3624 | NDB | 'Diagonal band nucleus' | 'Pallidum' |
| 102 | 0,3539 | SMT | 'Submedial nucleus of the thalamus' | 'Thalamus' |
| 103 | 0,3449 | LP | 'Lateral posterior nucleus of the thalamus' | 'Thalamus' |
| 104 | 0,3309 | PRNr | 'Pontine reticular nucleus' | 'Pons' |
| 105 | 0,3304 | SCm | 'Superior colliculus, motor related' | 'Midbrain' |
| 106 | 0,3281 | VM | 'Ventral medial nucleus of the thalamus' | 'Thalamus' |
| 107 | 0,3258 | SUT | 'Supratrigeminal nucleus' | 'Pons' |
| 108 | 0,3229 | VPM | 'Ventral posteromedial nucleus of the thalamus' | 'Thalamus' |
| 109 | 0,3194 | MA | 'Magnocellular nucleus' | 'Pallidum' |
| 110 | 0,2988 | SI | 'Substantia innominata' | 'Pallidum' |
| 111 | 0,2948 | LGd | 'Dorsal part of the lateral geniculate complex' | 'Thalamus' |
| 112 | 0,2895 | RT | 'Reticular nucleus of the thalamus' | 'Thalamus' |
| 113 | 0,2835 | PA | 'Posterior amygdalar nucleus' | 'Cortical Subplate' |
| 114 | 0,2746 | OT | 'Olfactory tubercle' | 'Striatum' |
| 115 | 0,2700 | CA3 | 'Field CA3' | 'Hippocampal Formation' |
| 116 | 0,2678 | LSr | 'Lateral septal nucleus, rostral (rostroventral) part' | 'Striatum' |
| 117 | 0,2649 | PAG | 'Periaqueductal gray' | 'Midbrain' |
| 118 | 0,2612 | PF | 'Parafascicular nucleus' | 'Thalamus' |
| 119 | 0,2605 | ICd | 'Inferior colliculus, dorsal nucleus' | 'Midbrain' |
| 120 | 0,2564 | SNc | 'Substantia nigra, compact part' | 'Midbrain' |
| 121 | 0,2529 | BST | 'Bed nuclei of the stria terminalis' | 'Pallidum' |
| 122 | 0,2521 | MH | 'Medial habenula' | 'Thalamus' |
| 123 | 0,2474 | SPFm | 'Subparafascicular nucleus, magnocellular part' | 'Thalamus' |
| 124 | 0,2417 | STN | 'Subthalamic nucleus' | 'Hypothalamus' |
| 125 | 0,2370 | VPMpc | 'Ventral posteromedial nucleus of the thalamus, parvicellular part' | 'Thalamus' |
| 126 | 0,2357 | LHA | 'Lateral hypothalamic area' | 'Hypothalamus' |
| 127 | 0,2292 | TR | 'Postpiriform transition area' | 'Olfactory Areas' |
| 128 | 0,2174 | ACB | 'Nucleus accumbens' | 'Striatum' |
| 129 | 0,2105 | LSc | 'Lateral septal nucleus, caudal (caudodorsal) part' | 'Striatum' |
| 130 | 0,2064 | ENTm | 'Entorhinal area, medial part, dorsal zone' | 'Hippocampal Formation' |
| 131 | 0,2061 | V | 'Motor nucleus of trigeminal' | 'Pons' |
| 132 | 0,2048 | CA1 | 'Field CA1' | 'Hippocampal Formation' |
| 133 | 0,1999 | GPi | 'Globus pallidus, internal segment' | 'Pallidum' |
| 134 | 0,1956 | PERI | 'Perirhinal area' | 'Isocortex' |
| 135 | 0,1856 | PMd | 'Dorsal premammillary nucleus' | 'Hypothalamus' |
| 136 | 0,1842 | AHN | 'Anterior hypothalamic nucleus' | 'Hypothalamus' |
| 137 | 0,1826 | MPN | 'Medial preoptic nucleus' | 'Hypothalamus' |
| 138 | 0,1809 | TRN | 'Tegmental reticular nucleus' | 'Pons' |
| 139 | 0,1807 | VTA | 'Ventral tegmental area' | 'Midbrain' |
| 140 | 0,1637 | PSV | 'Principal sensory nucleus of the trigeminal' | 'Pons' |

| Ranking | Node Modulation Index (Cohen's D) | Short Name | Full Name | Structure |
|---|---|---|---|---|
| 141 | 0,1571 | LSv | 'Lateral septal nucleus, ventral part' | 'Striatum' |
| 142 | 0,1525 | MPO | 'Medial preoptic area' | 'Hypothalamus' |
| 143 | 0,1491 | SPA | 'Subparafascicular area' | 'Thalamus' |
| 144 | 0,1416 | MRN | 'Midbrain reticular nucleus' | 'Midbrain' |
| 145 | 0,1415 | VMH | 'Ventromedial hypothalamic nucleus' | 'Hypothalamus' |
| 146 | 0,1340 | NI | 'Nucleus incertus' | 'Pons' |
| 147 | 0,1212 | NLL | 'Nucleus of the lateral lemniscus' | 'Pons' |
| 148 | 0,1172 | PRE | 'Presubiculum' | 'Hippocampal Formation' |
| 149 | 0,1155 | COAp | 'Cortical amygdalar area, posterior part' | 'Olfactory Areas' |
| 150 | 0,0950 | MGm | 'Medial geniculate complex, medial part' | 'Thalamus' |
| 151 | 0,0748 | VPL | 'Ventral posterolateral nucleus of the thalamus' | 'Thalamus' |
| 152 | 0,0288 | CA2 | 'Field CA2' | 'Hippocampal Formation' |
| 153 | 0,0269 | RR | 'Midbrain reticular nucleus, retrorubral area' | 'Midbrain' |
| 154 | 0,0134 | POL | 'Posterior limiting nucleus of the thalamus' | 'Thalamus' |
| 155 | 0,0092 | APN | 'Anterior pretectal nucleus' | 'Midbrain' |
| 156 | -0,0014 | LPO | 'Lateral preoptic area' | 'Hypothalamus' |
| 157 | -0,0048 | PAR | 'Parasubiculum' | 'Hippocampal Formation' |
| 158 | -0,0055 | SBPV | 'Subparaventricular zone' | 'Hypothalamus' |
| 159 | -0,0638 | RN | 'Red nucleus' | 'Midbrain' |
| 160 | -0,0854 | NPC | 'Nucleus of the posterior commissure' | 'Midbrain' |
| 161 | -0,2163 | PVp | 'Periventricular hypothalamic nucleus, posterior part' | 'Hypothalamus' |
| 162 | -0,2176 | RCH | 'Retrochiasmatic area' | 'Hypothalamus' |
| 163 | -0,2413 | ARH | 'Arcuate hypothalamic nucleus' | 'Hypothalamus' |
| 164 | -0,4835 | SPFp | 'Subparafascicular nucleus, parvicellular part' | 'Thalamus' |
| 165 | -0,5394 | TU | 'Tuberal nucleus' | 'Hypothalamus' |
